# Deep learning methods for designing proteins scaffolding functional sites

**DOI:** 10.1101/2021.11.10.468128

**Authors:** Jue Wang, Sidney Lisanza, David Juergens, Doug Tischer, Ivan Anishchenko, Minkyung Baek, Joseph L. Watson, Jung Ho Chun, Lukas F. Milles, Justas Dauparas, Marc Expòsit, Wei Yang, Amijai Saragovi, Sergey Ovchinnikov, David Baker

## Abstract

Current approaches to *de novo* design of proteins harboring a desired binding or catalytic motif require pre-specification of an overall fold or secondary structure composition, and hence considerable trial and error can be required to identify protein structures capable of scaffolding an arbitrary functional site. Here we describe two complementary approaches to the general functional site design problem that employ the RosettaFold and AlphaFold neural networks which map input sequences to predicted structures. In the first “constrained hallucination” approach, we carry out gradient descent in sequence space to optimize a loss function which simultaneously rewards recapitulation of the desired functional site and the ideality of the surrounding scaffold, supplemented with problem-specific interaction terms, to design candidate immunogens presenting epitopes recognized by neutralizing antibodies, receptor traps for escape-resistant viral inhibition, metalloproteins and enzymes, and target binding proteins with designed interfaces expanding around known binding motifs. In the second “missing information recovery” approach, we start from the desired functional site and jointly fill in the missing sequence and structure information needed to complete the protein in a single forward pass through an updated RoseTTAFold trained to recover sequence from structure in addition to structure from sequence. We show that the two approaches have considerable synergy, and AlphaFold2 structure prediction calculations suggest that the approaches can accurately generate proteins containing a very wide array of functional sites.

## Main text

The biochemical functions of proteins are generally carried out by a small number of residues in a protein which constitute a functional site--for example, an enzyme active site or a protein or small molecule binding site--and hence the design of proteins with new functions can be divided into two steps. The first step is to identify functional site geometries and amino acid identities which produce the desired activity--this can be done using quantum chemistry calculations in the enzyme case (to identify ideal theozymes for catalyzing a desired reaction) (*1*– *3*) or fragment docking calculations in the protein binder case (*4, 5*); alternatively functional sites can be extracted from native protein having the desired activity (*6, 7*). In this paper, we focus on the second step: given a functional site description from any source, design an amino acid sequence which folds up to a three dimensional structure containing the site. Methods have been developed for functional site scaffolding for sites made up of one or two contiguous chain segments (*6*– *10*), but with the exception of helical bundles (*8*) these do not extend readily to more complex sites composed of three or more chain segments. Current methods also have the limitations that assumptions must be made about the secondary structure of the scaffold, and that the amino acid sequence must be generated in a subsequent sequence step, so there is no guarantee that the generated backbones are in fact designable (encodable by some amino acid sequence).

An ideal method for functional de novo protein design would 1) embed the functional site with minimal distortion in a designable scaffold protein; 2) be applicable to arbitrary site geometries, searching over all possible scaffold topologies and secondary structure compositions for those optimal for harboring the specified site, and 3) jointly generate backbone structure and amino acid sequence. We reasoned that the trRosetta neural network (*11*), which maps input sequences to predicted structures, could be adapted for this purpose. Completely new proteins can be designed using trRosetta by starting from a random amino acid sequence, and carrying out Monte Carlo sampling in sequence space maximizing the probability that the sequence folds to some (unspecified) three dimensional structure (*12*). We refer to this process as “hallucination” as it produces solutions that the network considers ideal proteins but do not correspond to any actual natural protein (Fig. 1A); crystal and NMR structures confirm that the hallucinated sequences fold to the hallucinated structures (*12*). trRosetta can also be used to design sequences that fold into a target backbone structure by carrying out sequence optimization using a structure recapitulation loss function that rewards similarity of the predicted structure to the target structure (*13*). We sought to extend this approach to scaffold functional sites using trRosetta by sampling in sequence space with a combination of the hallucination loss to favor folding to a unique structure, and a structure recapitulation loss to favor formation of the desired functional site (rather than the entire structure as in (*13*); Fig. 1B; Methods). While we succeeded in generating structures that had segments which closely recapitulated functional sites, Rosetta structure predictions suggested that the sequences poorly encoded the structures, and hence we used Rosetta design calculations to generate more optimal sequences (*14*). Several designs targeting PD-L1 generated by constrained hallucination with binding motifs derived from PD-1, followed by Rosetta design, were found to have binding affinities in the mid-nanomolar range (Fig. S1). While this experimental validation is encouraging, the requirement for sequence design using Rosetta is at odds with property (3) above-the joint design of sequence and structure.

**Figure 1.**
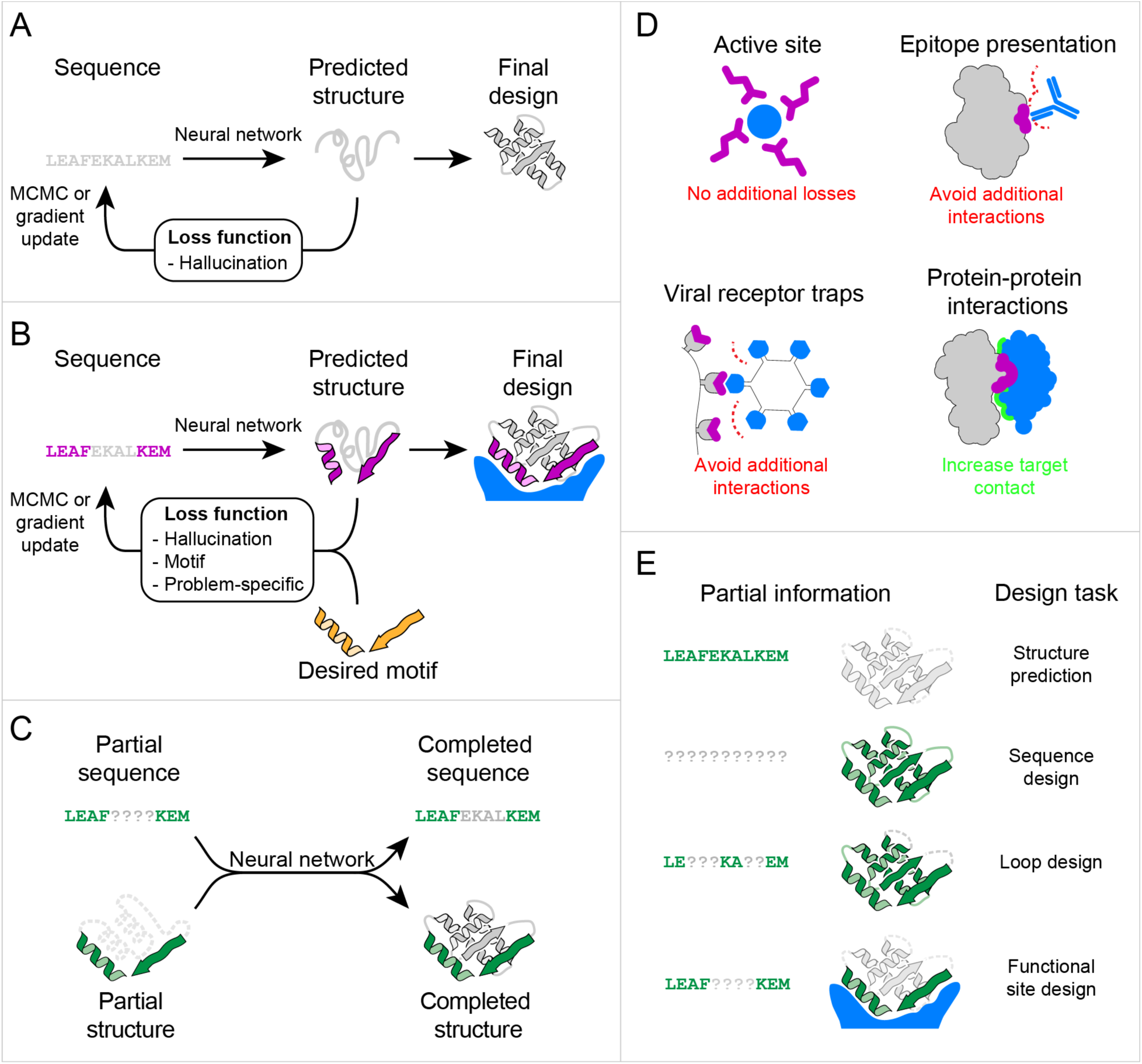
Methods for protein function design. (A) Free hallucination. At each iteration, a sequence is passed to the trRosetta or RoseTTAFold neural network, which predicts 3D coordinates and residue-residue distances and orientations (Fig. S3) which are scored by a loss function that rewards certainty of the predicted structure. The sequence is updated either by back propagating the gradient of the loss to the inputs or by MCMC, and passed back into the network for the next iteration. (B) Constrained hallucination. Same approach as in (A) but the loss function rewards motif recapitulation and other task-specific functions in addition to structural certainty. (C) Missing information recovery. Partial sequence and/or structural information is input into the network, and complete sequence and structure are output. (D) Design problems that can be addressed by constrained hallucination, and the corresponding loss functions (Fig. S3; Methods). (E) Protein design challenges formulated as missing information recovery problems. Colors in all panels: native functional motif (orange); hallucinated scaffold (gray); constrained motif (purple); binding partner (blue); non-masked region (green); masked region (light gray, dotted lines)

We found following the development of RosettaFold (*15*) that using it, rather than trRosetta, to guide motif-constrained hallucination resulted in designed protein sequences that more strongly encoded their structures (Fig. S2), likely reflecting the better overall modeling of protein sequence-structure relationships evidenced by the superior structure prediction performance (*15*). Constrained hallucination with RosettaFold has the further advantages that since 3D coordinates are explicitly modeled (trRosetta only generates residue-residue distances and orientations), motif recapitulation can be assessed at the coordinate level, and additional problem-specific loss terms can be implemented in coordinate space that assess interactions with a protein target (Fig. 1B, 1D).

In the following sections, we explore the use of the constrained RosettaFold hallucination method to design proteins containing a wide range of functionally diverse motifs (Fig. 2-4, Table S1). It is impractical to experimentally validate many designs for many different applications; we instead evaluate these designs using the AlphaFold (AF) protein structure prediction network (*16*) which has very high accuracy on *de novo* designed proteins (*17*). Although RoseTTAFold was inspired by AF, the two models were developed and trained independently, and hence AF predictions can be regarded as an orthogonal *in silico* test of whether RF designed sequences fold into the intended structures, analogous to traditional *ab initio* folding benchmarks (*13, 18*). For almost all problems, we obtained designs that are closely recapitulated by AF with overall and motif RMSD typically <2 Å and <1 Å respectively with model confidence pLDDT > 80 (Table S2). While solving current challenges with protein design clearly requires making and characterizing proteins in the lab, this *in silico* AF test is well suited for testing performance of design methods on a wide range of problems, and is quite stringent, as discussed below.

**Figure 2.**
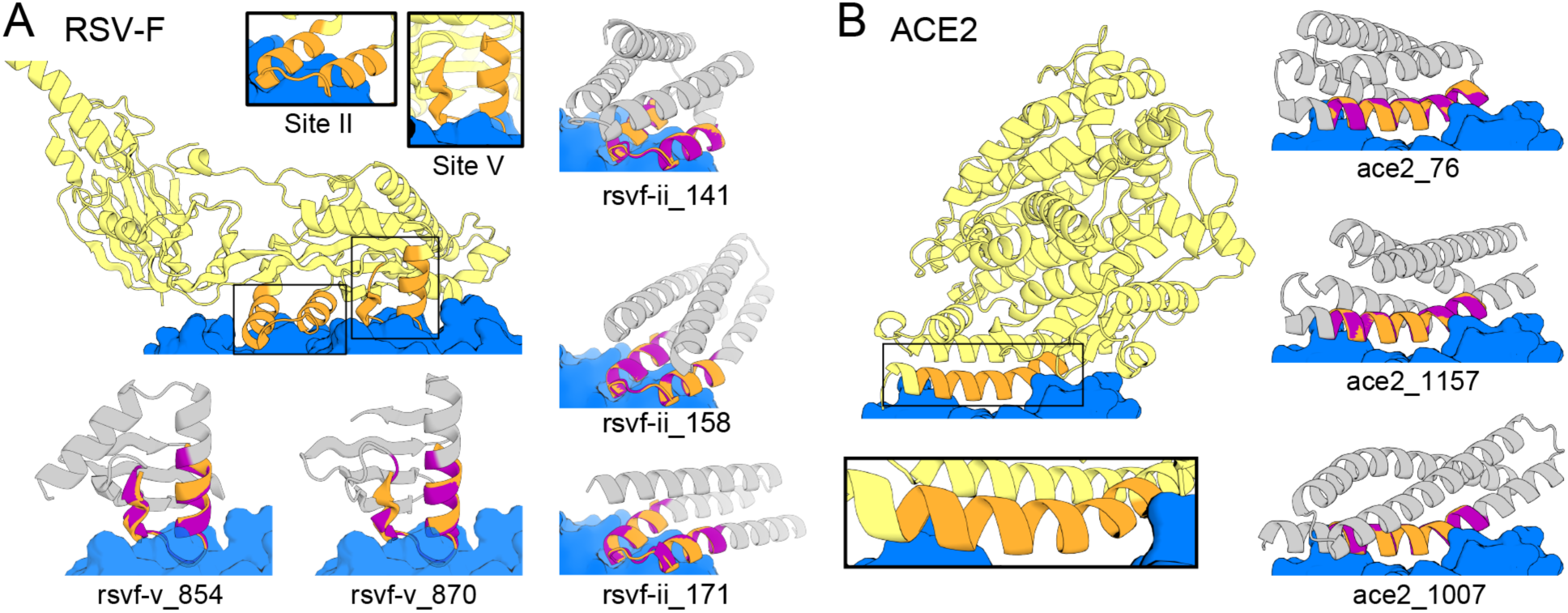
Hallucination of epitope scaffolds and receptor traps. (A) Design of proteins scaffolding immunogenic epitopes on RSV protein F (site II: PDB 3IXT chain P residues 254-277; site V: 5TPN chain A residues 163-181). Comparisons of the RF hallucinated models to unbiased AF2 structure predictions from the design sequence are in Fig. S8; here because of space constraints we show only the AF2 model; the two are very close in all cases. Here and in the following figures, we assess the extent of success in designing sequences which fold to structures harboring the desired motif through two metrics computed on the AF2 predictions: prediction confidence (AF pLDDT), and the accuracy of recapitulation of the original scaffolded motif (motif RMSD AF versus native). For RSV-F designs, these metrics are rsvf_ii_141 (85.0, 0.53 Å), rsvf_ii_158 (82.9, 0.51 Å), rsvf_ii_171 (88.4, 0.69 Å); rsvf-v_854 (81.5, 0.75 Å); rsvf-v_870 (80.4, 0.76 Å). (B) Design of COVID-19 receptor trap based on ACE2 interface helix (6VW1 chain A residues 24-42). Design metrics: ace2_76 (89.1, 0.55 Å); ace2_1157 (80.4, 0.47 Å); ace2_1007 (83.3, 0.57 Å). Colors: native protein scaffold (light yellow); native functional motif (orange); hallucinated scaffold (gray); hallucinated motif (purple); binding partner (blue). See Table S2 for additional metrics on each design.

### Hallucinating immunogen candidates and receptor traps

We first applied the constrained hallucination method to the problem of antigen presentation for immunogen design, where the goal is to scaffold a native epitope recognized by a neutralizing antibody as accurately as possible (and thus elicit antibodies binding the target protein upon immunization). Additional interactions with the target antibody are undesirable because the goal is to elicit antibodies recognizing the original antigen, and hence we incorporate an additional repulsive term assessed on the complex 3D coordinates in the composite loss function to penalize interactions with the antibody beyond those present in the epitope being scaffolded (Fig. 1D, S3). As a test case, we focused on respiratory syncytial virus, a leading cause of infant mortality whose F protein (RSV-F) contains antigenic epitopes for which structures with neutralizing antibodies have been determined (*7, 9, 10*). We sought to scaffold RSV-F site II, a contiguous helix-turn-helix motif that had previously been grafted successfully onto a 3-helix bundle architecture (*7*), as well as RSV-F site V, a helix-turn-strand motif that has not yet been scaffolded successfully (*19*). We were able to hallucinate designs for both epitopes with a variety of folds and motifs recapitulated to sub-angstrom C*α* RMSD in the AF predicted structure of the designed sequence (Fig. 2A, Fig. S8, S11; for these and all designs below, full amino acid sequence and PDB files are in the SM, and comparisons of the design models to AF predictions, in Fig. S8-10--since they are virtually identical, to save space we show only one of these in the main text figures).

We next applied the hallucination method to the design of receptor traps, which neutralize viruses by mimicking their natural binding targets and thus are inherently robust against mutational escape. We again augmented the loss function with an explicit penalty on interactions beyond those present in the receptor to avoid opportunities for viral escape. As a test case, we scaffolded the interfacial helix of human angiotensin-converting enzyme 2 (hACE2) interacting with the receptor-binding domain (RBD) of severe acute respiratory syndrome coronavirus 2 (SARS-CoV-2) spike protein (*20*). The hallucinated hACE2 mimetics have a diverse set of helical topologies, and AF2 structure predictions recapitulate the binding interface with sub Å accuracy (Fig. 2B, S8, S10).

### Hallucinating metal binding and enzyme active sites

We next explored the scaffolding of functional sites involved in metal-binding and catalysis. We designed scaffolds around a di-iron binding site, which is important in biological systems for iron storage (*21*) and also potentially harnessable for catalysis (*22, 23*). The motif, composed of four roughly parallel helical segments from *E. coli* bacterioferritin (cytochrome b1), was recapitulated with sub-angstrom RMSDs (Fig. 3A), in scaffolds with quite different helix connectivities than the parent (Fig. S9). For the calcium-binding EF-hand motif (*24*) composed of a 12 residue loop flanked by helices, the hallucination method readily generates a variety of scaffolds recapitulating either 1 or 2 EF-hand motifs within 0.5 Å RMSD of the calcium binding motif (Fig. 3C). When tasked with scaffolding one EF-hand motif, the method chooses to buttress the loop with a helix, avoiding the need for another long loop.

**Figure 3.**
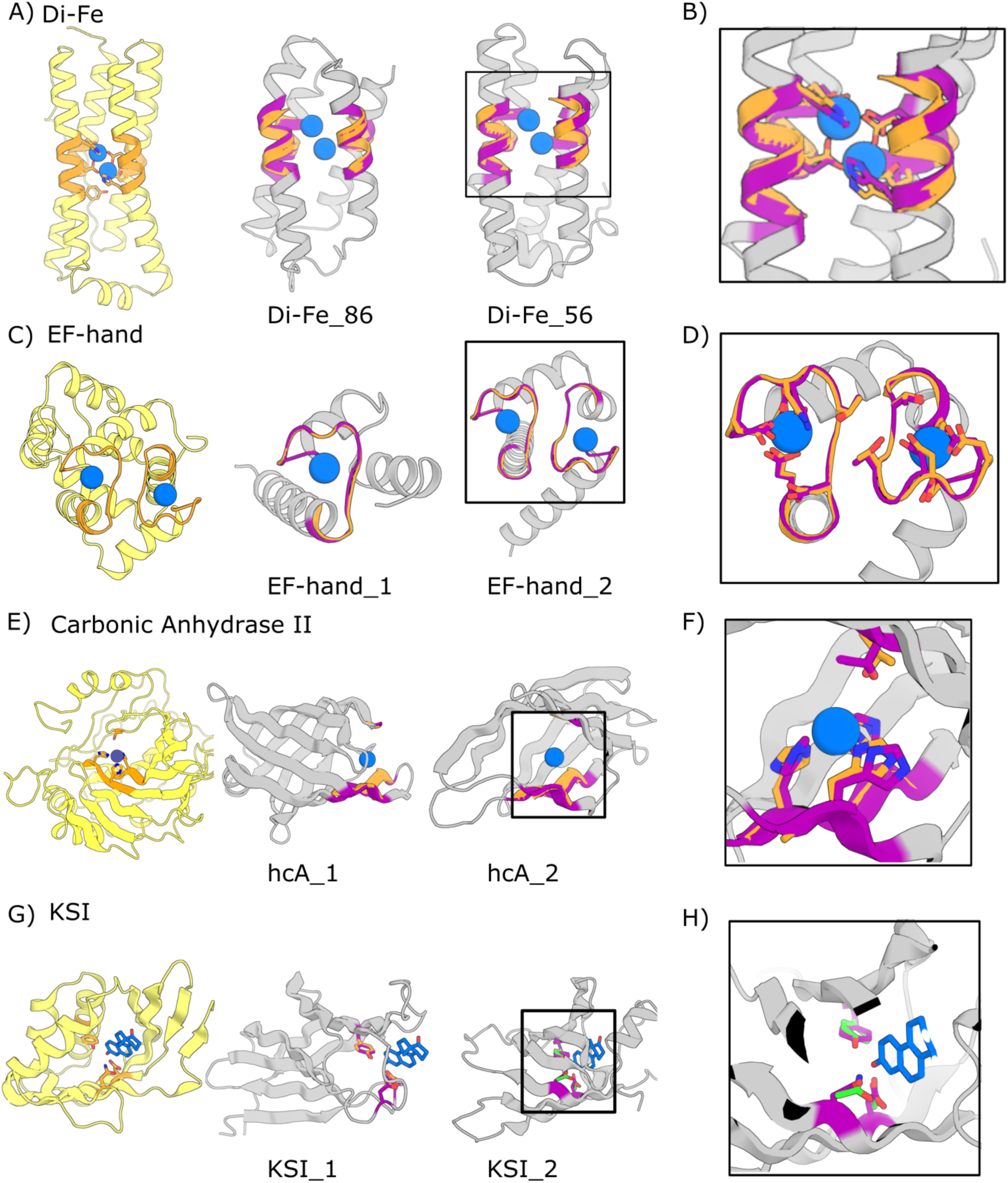
Hallucination of metal binding and enzyme active sites. (A-F) Hallucinations using backbone description of site using RF. (G-H) Hallucination using sidechain description of site using trRosetta followed by AF2. (A) Di-iron binding site from E. coli cytochrome b1 (1BCF chain A residues 18-25, 27-54, 94-97, 123-130). (C) EF-hand Calcium binding site. (E) Carbonic anhydrase II active site (5YUI chain A residues 62-65, 93-97, 118-120). (G) Δ^5^-3-ketosteroid Isomerase active site (1QJG chain A residues 14, 38, 99). Colors: native protein scaffold (light yellow); native functional motif (orange); hallucinated scaffold (gray); hallucinated motif (purple); bound metal (blue). Active site residues shown for boxed designs in panel B, D, F, and H for di-iron, EF-hand, carbonic anhydrase II, and Δ^5^-3-Ketosteroid Isomerase respectively. Design metrics (AF pLDDT, motif RMSD AF versus native): Di-Fe_86 (84, 0.90 Å), Di-Fe_56 (84, 0.86 Å) EF-hand_1 (84, 0.37 Å), EF-hand_2 (80, 0.37 Å), hcA_1 (73, 1.04 Å), hcA_2 (71, 0.62 Å), KSI_1 (84, 0.30 Å Cb), KSI_2 (72, 0.53 Å Cb)

We next sought to hallucinate enzyme active sites. Carbonic anhydrase II, which catalyzes the interconversion of carbon dioxide and bicarbonate, enables CO2 transport in humans (*25*), plays a key role in photosynthesis (*26*), and is emerging as a tool for CO2 sequestration (*27*). The active site contains 3 Zn^2+^ coordinating histidines (PDB ID 5yui: His94,His96,His119) on two strands, and a hydrophobic loop containing Thr199 which sequesters and orients the CO2. Despite the complexity of the irregular, discontinuous, 3 segment site, the method generated designs with sub angstrom motif RMSDs with correct His placement for Zn^2+^ coordination (Fig. 3E, S9); these are less than 100 residues, significantly smaller than the 261 residue long native protein.

To enable specification of sidechain geometry, we carried out iterative gradient descent using gradient information obtained by backpropagation through the AF neural network rather than RF, which currently does not explicitly model side chains (see Methods). As a test, we used the catalytic sidechain geometry of Δ^5^-3-ketosteroid isomerase (1QJG: residues 14, 38, 99), which catalyzes the isomerization of Δ^5^- to Δ^4^-3-ketosteroid needed for synthesis of steroid hormones in mammals (*28*). In initial experiments, we were only able to obtain designs that fully recapitulated the catalytic sidechain geometry when optimization was over a multiple sequence alignment rather than a single sequence; the landscape may be too rugged with the high resolution sidechain-based loss in the single sequence case. To overcome this problem, we developed a two-stage approach; with a first stage using both AF and trRosetta (to reduce the structure-prediction resolution and thus smoothen the loss landscape) and a description of the active site at the backbone level, followed by a second all-atom AF-only stage once the overall backbone was roughly in place. This two-stage approach led to multiple plausible solutions with predicted structures having a nearly exact match to the catalytic sidechain geometry (Fig. 3G, S9); however, we cannot use AF as an independent test of design accuracy in this case (given the very large number of model parameters, direct optimization against the output of a neural network has the potential to identify false optima, and hence independent *in silico* validation is important).

### Hallucinating protein-protein interfaces

We next sought to design binding proteins which extend beyond an input binding motif to make additional favorable interactions with the target by explicitly including the sequence and structure of the target in the hallucination process (Figs S6, Methods). We designed binders of the anti-inflammatory cytokine interleukin 10 (IL-10) *α*-receptor that incorporate one of the two discontinuous binding sites in the domain-swapped IL10 dimer in a single chain; the resulting scaffolds recapitulate the IL10 binding region within 0.5A (Fig. 4A, S10). Starting from the complement cascade protein C3d which enhances immune responses to covalently attached antigens (*29*) we designed binders to complement receptor 2 (CR2) present on B-cell and dendritic cells (*30*). The designs are much smaller (<100 AAs) than native C3d (306 AAs), recapitulate the binding interface with sub angstrom accuracy (Fig. 4B, S6C).

**Figure 4.**
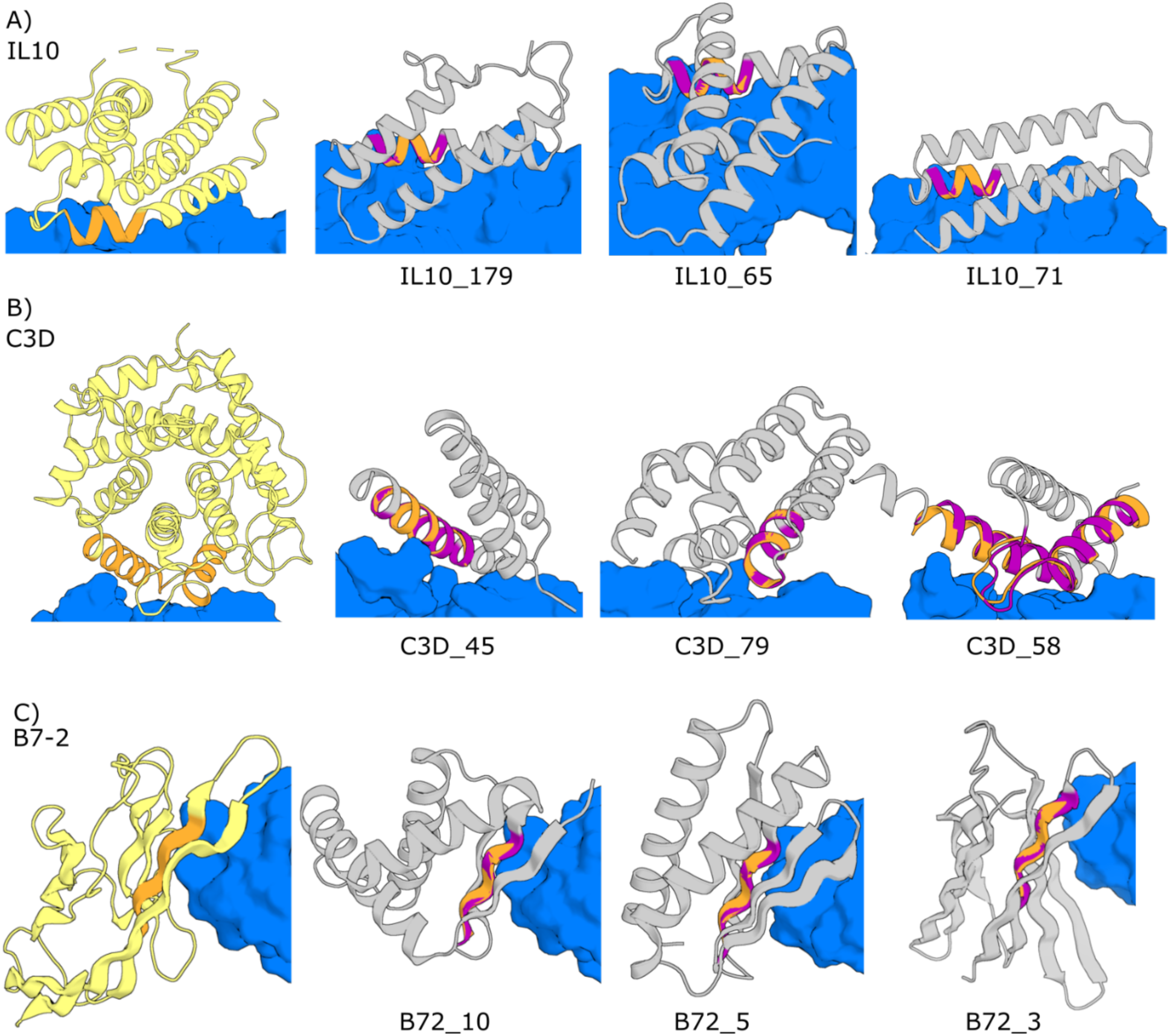
Hallucination of protein-protein interactions. Designs containing extended target binding interfaces built around native complex derived binding motifs. Targets are in blue and native scaffolds in yellow. (A) Target: IL10 receptor; scaffold: Interleukin 10 (1Y6K chain A residues 23-29). (B) Target: complement receptor; scaffold: Complement protein C3d (1GHQ chain A 104-126, 170-185). (C) B7-2 (1I85 chain B residues 84-88). Native functional motifs (orange); hallucinated scaffold (gray); hallucinated motif (purple). Design metrics (AF pLDDT, motif RMSD AF versus native): IL10_179 (82, 0.35 Å), IL10_65 (88, 0.37 Å), IL10_71 (75, 0.45 Å), C3D_45 (81, 0.71 Å), C3D_79 (70, 0.28 Å), C3D_58 (86, 0.47 Å), B72_10 (81, 0.29 Å), B72_5 (87, 0.23 Å), B72_3 (81, 0.25 Å)

As a test of building around beta strand motifs, we sought to design binders of the immune checkpoint protein CTLA-4 starting from B7-2, which binds CTLA-4 through four beta strands. Starting from a single five residue strand, hallucination in the presence of CTLA-4 generated designs having both alpha-beta and all beta topologies with novel binding modes and comparable interface contacts to native B7-2 (Fig. 4C, S10). As expected, designs hallucinated in the presence of the target had considerably better Rosetta protein-protein interface metrics (*4*) (binding free energy, etc) than those designed without the receptor (Fig. S6).

### Generalized protein function design by missing information recovery using RoseTTAFold

While quite powerful and general, the constrained hallucination approach is compute intensive, as a forward and backward pass through the network is required for each gradient descent step in sequence optimization. In the original training of RosettaFold for structure prediction a small fraction (15%) of tokens in the MSA are masked, and the network learns to recover this missing sequence information in addition to predicting structure. We reasoned that this ability to recover sequence information along with structural information could provide a second solution to the functional site scaffolding problem: given a functional site description, a forward pass through the network could potentially be used to complete, or “inpaint”, both protein sequence and structure (Fig. 1C; Methods). Here, the design challenge is formulated as an information recovery problem, analogous to the completion of a sentence given its first few words using language models (*31*) and completion of corrupted images using inpainting methods (*32*). As illustrated in Fig. 1E, a wide variety of protein structure prediction and design challenges can be similarly formulated as missing information recovery problems. We began from a RoseTTAFold model trained for structure prediction (*15*) and carried out further training on both fixed-backbone sequence design and fixed-sequence structure prediction tasks (Methods; Fig. S13; Algorithm S1). After training, the mean amino acid sequence recovery of the resulting model, denoted RFjoint, on a *de novo* protein test set was 33% (Fig. 5A; this is similar to Rosetta fixed backbone design performance), and there was also a slight increase in structure prediction accuracy (Fig. 5B). Thus, the model can both recover missing structure information given sequence and missing sequence information given structure.

**Figure 5.**
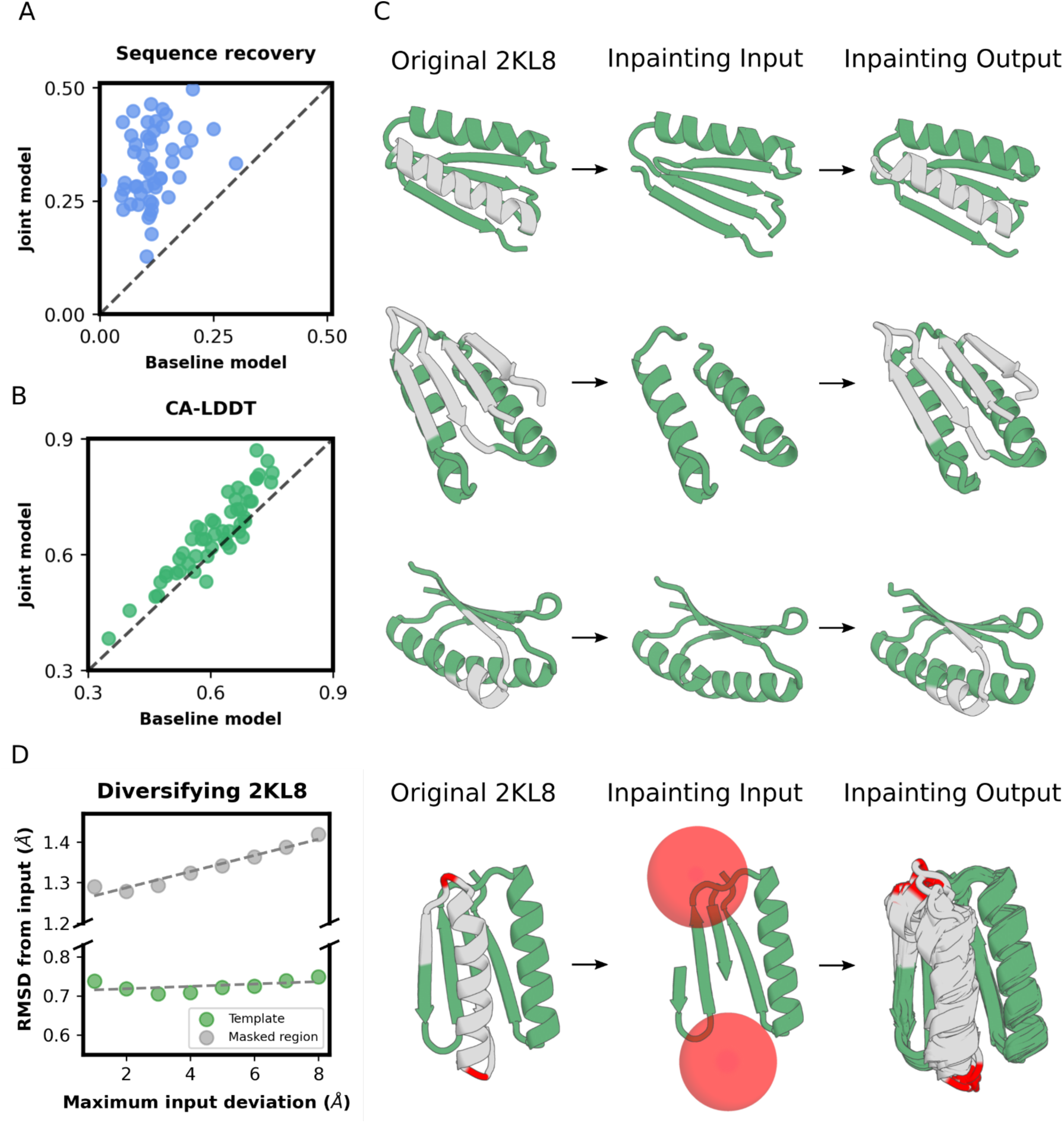
Joint sequence-structure recovery using RosettaFold. (A) Joint RoseTTAFold (RF*joint*) outperforms baseline RF in fixed-backbone sequence design on a held out set of *de novo* designed proteins. (B) RF*joint* preserves or exceeds the baseline model structure prediction quality on the *de novo* protein set. (C) Given a template sequence and structure (green) with regions of both sequence and structure masked (gray), RF*joint* can recover the missing sequence and structure in a single forward pass. The sequence and structure in contiguous regions of test set protein 2KL8 were both masked prior to input into RF*joint*. Top row: alpha helix. Middle row: four strand beta sheet. Bottom row: a 10-residue loop. (D) RF*joint2* builds sequence/structure between two given residue coordinates which enables tunable diversification of rebuilt segments. The depicted gray region was masked from 2KL8, and the two coordinates shown in red were randomly translated up to 8Å in any direction (within the illustrated red spheres). RF*joint2* is able to build back an ensemble of helical inpainted regions (right panel, AF2 predictions, AF2 pLDDT > 0.8 for all designs shown). Increasing structural diversity could be achieved in the central inpainted region (in both the RF inpainted structure models and the AF2 structure predictions of the inpainted sequences) by increasing the distance by which the red coordinates could be translated (left graph, gray points) without substantial disruption to the remainder of the template structure (left graph, green points, n=5000 structures/point).

We next considered design challenges where both sequence and structure information were missing for a portion of the protein. For smaller masked regions, the sequences and structures recovered by RFjoint are close to those of the input native structure, and as the size of the masked regions increases the divergence of both sequence and structure increases as expected (Fig. S14). The extent of variation in the resulting designs can be controlled by the amount of input sequence and structure information provided (Fig. S18C). Since the calculations require a single forward pass (including recycling outputs back as input) through the network, only 1-10 seconds on an NVIDIA RTX2080 GPU (Methods) are required to generate both sequence and structure.

Encouraged by the excellent performance of RF*joint* on simultaneous sequence and structure recovery despite being only trained on recovery of one or the other, we sought to improve this further by explicitly training on joint sequence/structure recovery tasks. Sequence and structure diversity is useful when designing proteins containing functional motifs, as subtle variations in the structure of the motif can drastically affect function (*33*), and hence we trained this new model to predict the sequence and structure of masked regions between two provided residue coordinates, in the absence of structural and sequence information of the residues flanking the two residue coordinates (to force the model to place structural elements based more on larger protein context than the local structure of the immediately connected chain segments). With this second model, which we call RFj*oint2*, the two residue coordinates can, at inference time, be varied, enabling the rapid generation of further sequence/structure diversity (Fig. 5D; a similar problem has been explored using Rosetta (*33*)). Of note, the degree of diversification in the inpainted region can be controlled by varying the distance by which the two residue coordinates are translated (Fig. 5D, left panel), while the structure of the templated (unmasked) protein remains remarkably stable.

We next explored the use of RF*joint* and RFj*oint2* to generate complete protein structures around the functional sites described in Figs 2-4, and found that success depended on the size and context of the input functional regi on. With the RF*joint* model, we found that best results were obtained for the more minimalist functional sites by first building up extended versions using the constrained hallucination approach. Many alternative structure and sequence completions can then be generated by RF*joint* in a network forward pass (Figure 6A, Figure S18). Almost all designs shown have sub-angstrom RMSD from the AF prediction to the native motif and <2 Å RMSD between design model and AF prediction (Fig. 6A, Fig. S19), and > 80 pLDDT. Diverse ensembles of such solutions to a specific design challenge can be very rapidly generated by varying the input sequence and structure information (Fig. S18). While RF*joint* struggled to generate well-predicted proteins from native/minimalist motifs, we found that RF*joint2* was able to generate complete and confidently-predicted (by AF2) protein models from smaller regions, such as a single EF hand motif (Fig. S18B). Further, RF*joint2* could simultaneously scaffold two motifs while retaining good (<1 Å RMSD) alignment to both (Fig. 6B, top row). Remarkably, in some cases, RF*joint2* was able to generate well-predicted scaffolds to complex, multi-chain motifs taken directly from a native crystal structure (Fig. 6B, middle and bottom row), as well as translationally symmetric proteins (Fig. S20), provided little more than the desired motif, in a single forward pass through the network.

**Figure 6.**
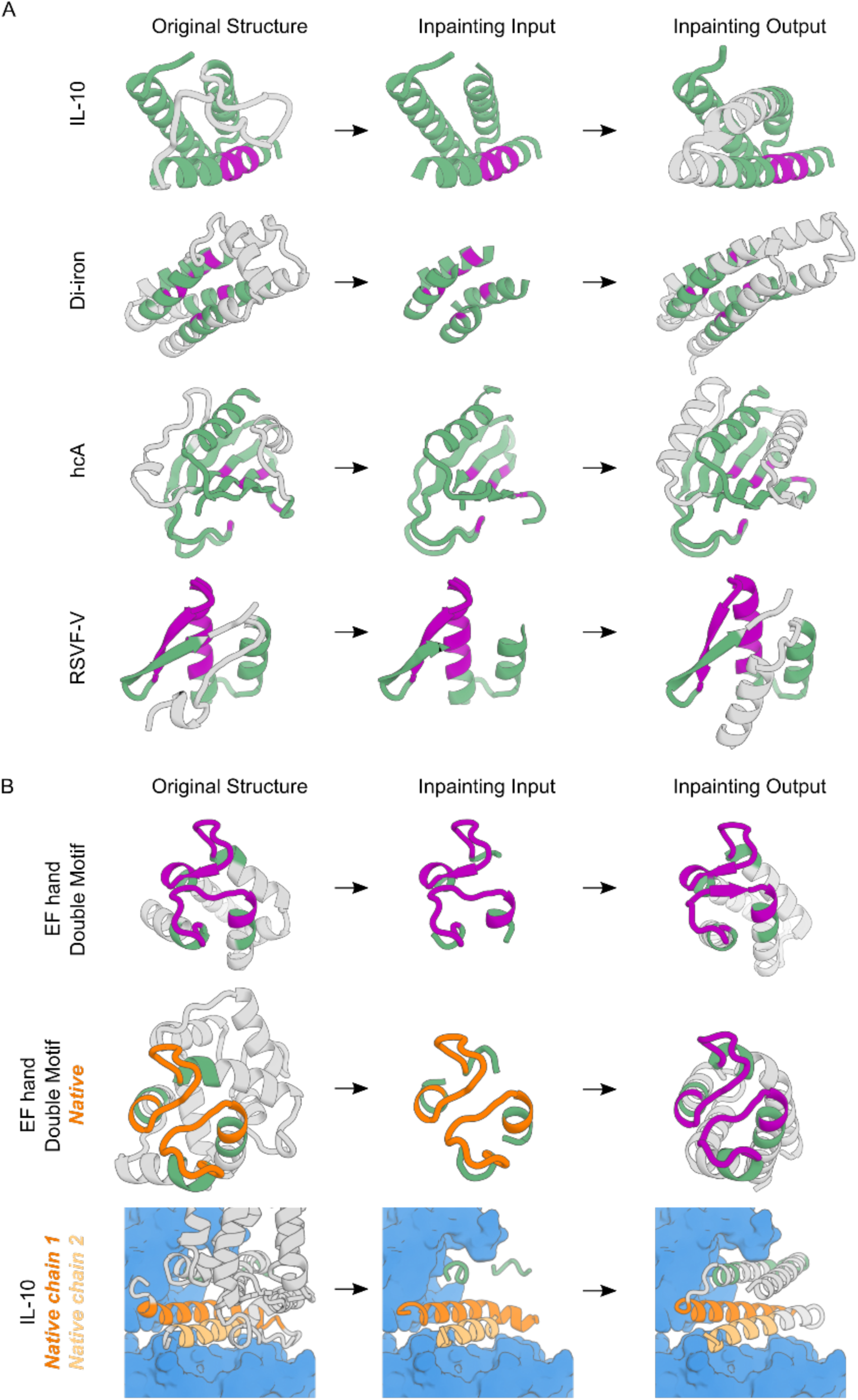
Protein function design by joint sequence-structure information recovery. Design of proteins harboring functional motifs via information recovery using RF*joint* and RF*joint2*. All structures of designs shown are the AF2 prediction of that design. In all cases, template inputs (sequence and structure) that are functional and their corresponding outputs are colored in purple, template inputs that are not directly related to function are in green, along with their corresponding outputs. Functional template inputs derived from a native structure are in orange, with corresponding outputs in purple. Depicted in gray are the regions of sequence and structure masked from the original protein (input column) or that were generated via RF*joint*/RF*joint2* (output column). (A) RF*joint* functional motif design examples. From top to bottom row with (AF2 motif RMSD to native, AF2 pLDDT): IL-10 (93.1, 0.57 Å), Di-Iron (91.0, 0.49 Å) carbonic anhydrase (78.8, 1.09 Å), RSVF-V (81.8, 1.39 Å). (B) RF*joint, 2* functional motif design examples. From top to bottom row with (AF pLDDT, motif RMSD AF vs native): EF hand double motif starting from a hallucination (85.4, 0.69 Å motif #1, 0.86 Å motif #2), EF hand double motif starting from native crystal structure (PDB: 1PRW, chain A 16-35, 52-71) (78.7, 1.13 Å motif #1, 1.10 Å motif #2), IL10 motif (light/dark orange) starting from native crystal structure (PDB: 6×93, chain A 16-41, 83-88, chain D 96-101,143-156) (75.6, 1.16 Å).

Tests on the full range of challenges described here suggest that the two function design approaches are complementary: the constrained hallucination approach can build protein structures harboring minimalist functional sites but is quite compute and memory intensive since it requires a forward and backward pass (to generate gradient information to guide sequence optimization) through the neural network at each step of sequence optimization (Methods), while the missing information recovery method in most but not all cases requires extended functional site description but is much less compute intensive, and generally outperforms the hallucination method when more starting information is provided, as illustrated by the lower RMSDs on constrained regions (Fig. S15). This difference in performance can be understood by considering the manifold in sequence-structure space corresponding to folded proteins; the space of all possible sequence-structure pairs is far larger than the set of sequence-structure pairs of folded proteins, and hence this manifold occupies a tiny fraction of the overall space. The missing information recovery approach can be viewed as projecting an incomplete or corrupted input sequence-structure pair onto the subset of this manifold (as represented by RosettaFold) containing the functional site--if insufficient starting information is provided, this projection is not necessarily well determined, but with sufficient information, it readily produces protein-like solutions, updating sequence and structure information simultaneously. The loss function used in the hallucination approach is constructed with the goal that minima lie in the protein manifold, but there will likely not be a perfect correspondence, and hence stochastic optimization of the loss function in sequence space may not produce as protein-like solutions as the inpainting approach-- on the other hand, since stochastic search can be initiated from any starting point, the hallucination approach can start from minimalist functional site descriptions, or, as in the fully unconstrained case (*12*), no sequence and structural information at all.

### Evaluation of designs using AF2

New protein design methods have traditionally been evaluated by experimental testing, and for actual applications it is essential to make and characterize proteins in the lab. The high structure prediction accuracy of AF2 now enables evaluation of new design methodology in silico, which has the considerable advantage that a much wider variety of design challenges can be evaluated. In the work described here, AF2 was not used for any of the design calculations except for the sidechain active site design case of Fig. 3E, and hence provides an independent test of design accuracy. Both the backbone design challenge--generating a plausible protein backbone with a geometry capable of hosting a desired site, and the sequence design challenge--generating a sequence which strongly encodes this backbone, are quite formidable. For the backbone design problem, the very large set of structures predicted for naturally occurring proteins using AF and recently made available (*34*) provides an excellent point of comparison: for the RSV-F site V immunogen design challenge described above, the frequency of non-homologous proteins in the AF proteomes database and the Protein Data Bank (PDB) (*35*) matching the functional site with equal or lower RMSDs than our designs was 3.9×10^−6^ (Fig. S17; Supplementary Text); similarly low frequencies of suitable natural scaffolds in the PDB were observed for other targets (Table S3). For the sequence design problem, the accuracy of native protein structure prediction based on single amino acid sequences provides a point of comparison; as shown in Fig. S16, our designs are predicted more confidently from sequence than the vast majority of native proteins with known crystal structures, and on par with structurally validated de novo designed proteins. This success in designing sequences confidently predicted to fold to structures harboring a wide range of functional sites derives in part from a key advance over classical protein design pipelines, which treat backbone generation and sequence design as two separate problems: our methods simultaneously generate both sequence and structure, taking advantage of the ability of RoseTTAFold to reason over and jointly optimize both data types.

## Conclusions

The deep learning methods presented here are quite general, requiring no inputs other than the structure and sequence of the desired functional site, and unlike current non-deep-learning methods, do not require specification of the secondary structure or topology of the scaffold, and simultaneously generate both sequence and structure. Despite a recent surge of interest in using machine learning to design protein sequences (*36*–*43*), the design of protein structure is relatively underexplored, likely due to the difficulty of efficiently representing and learning structure (*44*). Generative adversarial networks (GANs) and variational autoencoders (VAEs) trained on specific fold families have been used to design biophysically plausible protein backbones (*45, 46*), but not ones containing functional sites. RoseTTAFold and Alphafold have been trained on the entire PDB, and thus generalize from a very wide range of known protein structures. Our “activation maximization” hallucination approach enables use of arbitrary loss functions tailored to specific problems without retraining for any sequence length.

Complementary to this, the ability of our “missing information recovery” inpainting approach to expand from a given functional site to generate a coherent sequence-structure pair should find wide application in protein design because of its speed and generality. The combination of the two approaches is more powerful than either one alone, as ensembles of solutions to a given functional design problem can be generated very rapidly using the second approach starting from extended site descriptions identified in the first. The hallucination approach could, in theory, also be used to refine the more extensive designs generated by inpainting. The two approaches individually, and the combination of the two, should increase in power as more and more accurate protein structure, interface, and small molecule binding prediction networks are developed moving forward.

## Acknowledgements

We would like to thank Luki Goldschmidt for maintaining the computational resource in the IPD; Christoffer Norn for general discussions about trRosetta; Brian Coventry and Nathaniel Bennett for advice on interface design; Bruno Correia, Casper Goverde, and Karla Castro for advice on RSV-F epitopes and motif grafting methods; Ta-yi Yu, Gyu Rie Lee, Linna An, and Xinru Wang for advice on flow cytometry; Runze Dong and Varshan Muhunthan for exploratory analyses; Brian Trippe for feedback on the manuscript; Sam Pellock for expertise on enzyme design.

## Author contributions

Designed the research: JW, SL, DJ, DT, SO, DB

Developed the hallucination method: JW, SL, DT, IA, SO, JD

Developed the inpainting method: DJ, JLW, JW, DT, SL

Generated designs using hallucination: JW, SL, DT, SO

Generated designs using inpainting: DJ, JLW, JW, AS, SL

Analyzed data: JW, SL, DJ, DT, JLW, ME

Trained neural networks: DJ, MB, JLW

Performed experiments: JW, SL, LFM, JC, WY

Wrote the manuscript: JW, SL, DJ, DT, JLW, DB

## Data and materials availability

All code will be made publicly available upon publication.

## Supplementary materials

- Materials and Methods
- Supplementary Text
- Figures S1 - S21
- Tables S1 - S3
- Algorithm S1
- Data S1

## Data and code availability

All source code will be made freely available upon publication.

