## Supplementary materials for "Deep learning methods for designing proteins scaffolding functional sites"

This file contains:

- Materials and Methods
- Supplementary Text
- Figures S1 - S21
- Tables S1 - S3
- Algorithm S1
- Data S1

### Materials and Methods

#### Sequence representation

For structure prediction, the input to trRosetta and RoseTTAFold is a tensor $\mathrm{X} \in \mathbb{R}^{N \times L \times A}$ representing a one-hot-encoded multiple sequence alignment (MSA), where *L* is the sequence length, *N* is the number of aligned sequences, and $A = 21$ is the alphabet size (20 amino acids plus gap character, although gaps are never used during design). For design with RoseTTAFold, which was used for most of the designs in this paper, we optimized a single sequence ($N = 1$) and applied a 20% dropout, which is implemented at a variety of layers within the network. The PD-1 mimetics (Fig. S1) were designed using an earlier pipeline that used trRosetta and optimized a 1000-sequence MSA ($N = 1000$) with 0-20% dropout on input 2D features (*14*). Designing an MSA improves motif accuracy with trRosetta (*13*) but is not necessary when using RoseTTAFold. We find adding trRosetta loss during AlphaFold design to increase the number of designs that are within 1.5 RMSD and good starting points for semi-greedy search -- see details below.

#### Loss function

We optimize a loss function

$$\mathcal{L} = w_M \mathcal{L}_M + w_H \mathcal{L}_H + \mathcal{L}_{aux}$$

consisting of the motif loss $\mathcal{L}_M$, which scores the accuracy of the functional site in the design, and a hallucination loss $\mathcal{L}_H$, which scores how strongly the sequence encodes a backbone geometry (Fig. 1B), as well as optional auxiliary losses $\mathcal{L}_{aux}$ for specific tasks (Fig. S3; Supplementary Text). For all the designs in this paper we used $w_M = w_H = 1$.

For a protein of length *L*, the motif loss is defined as a negative cross-entropy between reference (one-hot-encoded) and predicted residue-residue geometric feature distributions $p(y)$:

$$\mathcal{L}_M = -\sum_{y\in\{d,\omega,\theta,\varphi,\theta^T,\varphi^T\}} \left[ \left( \sum_{i=1}^{L} \sum_{j\neq i}^{L} m_{ij} \, log \, p(y_{ij} = y_{ij}^0) \right) / \left( \sum_{i=1}^{L} \sum_{j\neq i}^{L} m_{ij} \right) \right]$$

where

$$m_{ij} = \begin{cases} 1, & \|C\beta_i - C\beta_j\| \le 20 \text{ and } i,j \in \text{motif} \\ 0, & \text{otherwise} \end{cases}$$

$y \in \{d, \omega, \theta, \varphi, \theta^T, \varphi^T\}$ represents residue-residue distances and orientation angles and $y^0$ is the value of the distance or angle in the reference motif. The features $d$ and $\omega$ are symmetric while the angles $\theta, \varphi$ are asymmetric, so $\theta^T$ and $\varphi^T$ are included to match the double-counting of $d$ and $\omega$ across the diagonal. This cross-entropy is averaged over all residue pairs in the motif, represented as a binary mask *m*. We restrict this loss to residue pairs within 20 Å because RoseTTAFold and trRosetta do not make quantitative predictions beyond this distance. In some cases we supplemented this cross-entropy motif loss with a backbone coordinate RMSD loss (Supplementary Text).

The hallucination loss is defined as the entropy of renormalized network predictions:

$$\mathcal{L}_H = \sum_{y\in\{d,\omega,\theta,\varphi,\theta^T,\varphi^T\}} \left[ \left( \sum_{i=1}^{L} \sum_{j\neq i}^{L} (1 - m_{ij}) H\big(\hat{p}(y_{ij})\big) \right) / \left( \sum_{i=1}^{L} \sum_{j\neq i}^{L} (1 - m_{ij}) \right) \right]$$

where the entropy over *k* geometric feature bins is defined as

$$H(p) = \sum_{k} p_k \log p_k$$

and $\hat{p}(y) = \exp(\beta \log p(y)) / \sum \exp(\beta \log p(y))$. Empirically, we found that performing this renormalization with $\beta = 10$, and only using bins up to 5 Å for the pairwise distance distributions $p(d)$, gave more realistic structures. In some cases we defined the hallucination loss using a KL divergence rather than entropy, which gave similar results (Fig. S3D; Supplementary Text).

#### Optimization methods

We used an MCMC method based on our previous work on unconstrained hallucination (*12*). Starting from a random sequence, single mutations were proposed and the loss function evaluated. The mutation was either accepted or rejected according to the standard Metropolis criterion. Acceptance temperature was 0.002 and annealed by exponential decay with a 500-step half-life; design quality was not sensitive to these parameters. For proteins around 120 residues long, we found this approach converged in about 30,000 steps and took about 90 minutes on Nvidia GeForce RTX2080 GPUs, which we used for all hallucination runs. Although slow, this approach has the advantage that mutations can include insertions and deletions, which is useful when redesigning loops.

We used a gradient-descent method based on our previous fixed backbone sequence design study (*13*). Starting with randomly initialized input logits $X \sim N_{N \times L \times A}(0, 0.01)$, we apply a softmax followed by an argmax operation to obtain a one-hot-encoding $X_{oh}$. To backpropagate the gradient of the loss $\nabla \mathcal{L}$ through the discrete one-hot sequence to the continuous logits, we employed a reparameterization trick(*13*, *47*) where gradients were passed through the one-hot sequence as if it had the softmax values of the logits (*48*, *49*). For a protein of length *L*, on optimization step *t*, we update the input logits with normalized gradients and a constant learning rate $\alpha$:

$$X^{(t+1)} \leftarrow X^{(t)} - \alpha \sqrt{L} \frac{\nabla \mathcal{L}}{||\nabla \mathcal{L}||}$$

Typically we used $\alpha = 0.05$, although results are reasonable for any $0.01 < \alpha < 0.2$(Fig. S4A). We also tested decaying the learning rate over time, but this did not outperform constant learning rate, as seen previously for fixed backbone hallucination (*13*). With trRosetta, we found that sampling from the softmax distribution over sequence logits (*47*) yielded higher DAN-lDDT and lower motif RMSD than simply taking the most probable sequence (argmax), but argmax was better when using RoseTTAFold.

Gradient-based optimization with trRosetta converged in 200 steps for a 120-residue protein, taking approximately 5 minutes on our GPUs, while RoseTTAFold took 400 steps or 10 minutes per design. A hybrid procedure of gradient descent followed by MCMC yielded slightly improved designs but required much more GPU time, while MCMC-only or MCMC followed by gradient descent yielded inferior results (Fig. S4B-C).

**Motif placement**

For most designs shown, at the beginning of optimization, each discontinuous segment of the motif is mapped to a random block of residue positions on the designed sequence. The motif loss is applied to these "constrained" regions, while the hallucination loss is applied to the remaining residue positions. The positions corresponding to the motif stay fixed during optimization. For each new problem, we start by specifying a range for the total protein length *L* and generate many designs with randomly sampled *L* from the range and randomly placed motif segments. We then identify the values of *L* and inter-segment gap lengths that yielded the best designs and run followup design trajectories with these parameters in order to deeply sample productive regions of the search space. We also developed algorithms which adaptively place motifs during optimization either by minimizing motif loss over all possible placements or performing a greedy search (Supplementary Text), but found that while useful for certain problems, these were not consistently better than the simpler fixed-placement strategy for the problems we tested (Fig. S5).

**Design of enzyme active sites using AlphaFold**

To design de novo scaffolds for the active site of $\Delta^5$-3-ketosteroid isomerase (*28*), we used AF in a two-stage method, the first stage focusing on backbone generation and the second on sidechain geometry optimization. In stage 1, we perform 200 steps of gradient descent to optimize a real-valued tensor $X \in \mathbb{R}^{1 \times L \times A}$ representing sequence logits. The argmax of the softmax of the logits is used as input to AF and trRosetta. To allow backprop through the argmax function, we use the gradient straight-through trick as described previously (*13*). Gradients are obtained from both AF and trRosetta, weighted equally, and used to update the logits X. Losses used for AF are the predicted LDDT and aligned error (for hallucination) and Cb distogram CCE (for motif recapitulation, defined similarly as the CCE used with RoseTTAFold above), sidechain FAPE (*16*) and RMSD (root-mean-squared-deviation); losses for trRosetta are KL divergence (Supplementary Text) and CCE, but excluding the theta dihedral. Stage 1 is run using the ADAM optimizer (*50*) with a learning rate of 5e-3. The gradients are normalized by the norm at each iteration. We found that if we do not use trRosetta as part of the loss, it is very difficult to achieve losses below 2 RMSD. In stage 2, the sequence from stage 1 is subjected to 400 steps of semi-greedy optimization using AF: at each step a random position is mutated, if the loss decreases, the mutation is accepted, if not, up to 20 independent random mutations are attempted. If none of the 20 mutations decreased loss, the mutation with best loss is accepted. For the first stage, 400 independent designs were generated. Each design had 3 random indices between 0 and 99 selected to define the positions of the active site. The top 4 designs were selected for stage 2. The loss for stage 2 is the weighted sum of predicted LDDT and aligned error, and sidechain FAPE and RMSD. The confidence loss was scaled by 0.01 and sidechain loss by 1.0. To attempt to avoid false local optima in a particular set of AF weights, during stage 2 we evaluated the loss using a randomly chosen one of 4 AF models (model_1_ptm, model_2_ptm, model_3_ptm, and model_5_ptm) (sets of weights) on each step. This is similar to averaging the 4 models (*51*, *52*) but is more compute efficient. We withhold model_4_ptm for validation -- the designs shown in the figures come from this model.

#### Motif selection

Because RoseTTAFold is most accurate when predicting secondary structures, we selected functional motifs with as little loop content as possible. For antigenic epitopes, viral receptor traps, and enzyme active sites, we chose the functional motifs based on previous structural literature. For binding interfaces, we identified interface residues as those within 5 Å of the binding partner and scaffolded motifs consisting of 2-4 contiguous blocks manually chosen to contain as many of the interface residues as possible. If the method struggled to produce designs accurately recapitulating the functional motif, we often found it helpful to include additional secondary structure elements buttressing the motif in the native structure. Table S1 lists the mimetic design targets, their PDB accessions, the residue numbers of constrained regions, and references.

#### Design selection

For each design problem, we generated between 200 and 2000 hallucinations (Fig. S7) and filtered them on AF pLDDT, motif RMSD of AF predictions, and (as extended unpacked helices often have high pLDDT) radius of gyration to 10-50 designs, and did a final manual visual filter to obtain the high-quality designs shown in the figures. The "model 4" weights were used for all AF predictions for filtering. This process comes with some risk of designing "adversarial examples", or sequence-structure pairs that score well by AF that do not fold or function in reality, due to the presence of artifactual minima in the loss landscape of the structure-prediction model (*53*, *54*). However, because we design using RoseTTAFold, which is trained independently of AF, any final designs must be well-predicted by two different networks, which is expected to act as a form of regularization and provide some robustness to adversarial examples. Supporting this idea, we find that our designs generated by RoseTTAFold and filtered by AF have similarly low variability in predicted structure and quality metrics between different neural networks as experimentally successful de novo proteins (Fig. S21). On the other hand, in our active-site designs where we directly optimize the sequence input to AlphaFold via gradient descent and MCMC, or in other recent works employing direct optimization against AF2 (sometimes using >10,000 AF predictions for a single design), we expect greater risk of adversarial examples (*51*, *52*). This is likely mitigated by using multiple AF models (*51*, *52*), but since the 5 AF models share early training checkpoints, they are likely less mutually orthogonal than RoseTTAFold and AF.

**Protein binder hallucination and interface refinement**

To design protein-binding proteins with interfaces expanded around their native interface motifs, our method took 2 stages. First, standard hallucination was done to scaffold a “stub”, or a small motif from the native interface, with repulsive and attractive losses active. This yields designs that are roughly shape-complementary to the binding partner (“target”), but does not have the ability to fine-tune contacts. To do this, we refined a small number of high-scoring designs by 100-1000 steps of MCMC with RoseTTAFold predicting the entire binder/target complex but only optimizing the binder sequence. We predicted complexes by concatenating the binder and target sequences with a 200 amino-acid gap between them in the residue index input to RoseTTAFold (*15*). RoseTTAFold has limited accuracy predicting native protein structures and complexes from single sequences. To ensure that the target is accurately predicted (as this is a prerequisite for accurately hallucinating interactions to it) we input the structure of the target plus the stub as homology templates to RoseTTAFold (Fig. S6). As expected, this usually yielded predictions of the target (and target-stub relative position) extremely close to the crystal structure. During 2-chain refinement, we only applied a motif loss to preserve the structure of the binder and its relative position to the target; no hallucination, repulsive, or attractive losses were used. Although in principle we could use gradient descent for joint binder/target optimization, or start from a random binder sequence rather than an existing binder hallucination, we found that MCMC refinement of a previously hallucinated monomer gave the best results.

**Training RoseTTAFold to jointly model sequence and structure**

Standard RoseTTAFold (*15*) (RF) has been trained on structure prediction (sequence inputs, structure outputs) using homolog templates (structure input). In the newer versions, we mask a portion of the input MSA and apply a loss to predictions of the masked amino acids (sequence output) to encourage the network to extract more meaning from the MSA (*16*, *55*). Starting with a pre-trained RoseTTAFold that has both sequence and structure inputs and outputs, we fine-tuned the model for an additional 5 epochs on both sequence design and structure prediction tasks with a learning rate of $5x10^{-4}$. In the fixed-backbone sequence design task, which comprised 75% of the fine-tuning examples, we replaced 90-100% of the sequence input with "mask" tokens while retaining the native backbone features as a template structure input (*15*). For the remaining 25% of fine-tuning examples, classic structure prediction was performed, with 15% of the MSA randomly masked, no masking of the query sequence, and inputting homolog template structures as usual (*15*). Note that all fine tuning examples used the same loss formula, which had increased weight on the cross entropy over sequence prediction logits (see Methods). We refer to this fine-tuned RoseTTAFold as $RF_{joint}$. Training curves for the sequence design and structure prediction tasks are shown in Fig. S13. As a control, we also started joint training from a completely untrained RoseTTAFold model, and saw training saturation at very poor losses on both sequence design and structure prediction (Fig. S13A). This suggests that pre-training on structure prediction was needed to achieve high performance on the sequence design task.

Through fine-tuning, $RF_{joint}$ *implicitly* learns to generate sequence and structure simultaneously when both are missing in a template input. We hypothesized that performance at this task could be further improved if it were *explicitly* trained (Fig. S13C). To explicitly train RoseTTAFold to simultaneously predict sequence and structure ($RF_{joint2}$), we started from the same pre-trained RoseTTAFold model. For the joint structure/sequence prediction task (50% of fine-tuning examples), contiguous regions of 10-25 amino acids comprising at least one full secondary structure element (helix, loop or strand) were masked out (Fig. S13D, gray). The sequence and structure of a further 3-5 'flanking' residues were masked out either side of this contiguous region (Fig. S13D, red). However, the distograms (but not angle maps or amino acid identity) were provided for the residue immediately N- and C-terminal to the central contiguous masked region (Fig. S13D, asterisks). Losses were applied only to the central masked region, with the flanking sequence/structure never revealed to the network. For the fixed backbone sequence design task (25% of fine-tuning examples), which was also included in the training, a similar strategy was used, with the network tasked with predicting the sequence of a masked region, naive to the sequence/structure of a 3-5 amino acid flanking region either side of the masked region. For the classic structure prediction task (25% of fine-tuning examples), 15% of the MSA was randomly masked, as described above.

**Joint sequence-structure inpainting with a jointly trained RoseTTAFold**

To apply $RF_{joint}$ to protein design, we input a sequence and structure, masking certain residues in the sequence by replacing them with mask tokens and masking corresponding residues in the structure by setting their template embeddings to zero (*15*). We then predict the structure and sequence logits for the entire protein. The output structure, including regions that were originally both masked and unmasked, is used as an updated design model, and the most probable predicted amino acid at each masked position (argmax) is taken to complete the sequence. We found that by recycling the predicted structure and inputting it again as a template without masking, along with the masked sequence, yielded improved outputs. Therefore, for most design problems, we ran 5-10 iterations of inpainting to get a final optimized design. If there is a functional motif whose structure must be maintained, we include the native structure of this motif as an additional template input. A single design using 10 iterations of inpainting takes 5 seconds on our GPUs.

The iterative inpainting method described above is deterministic. To sample ensembles of outputs with small variations in sequence and structure using $RF_{joint}$, we either vary the exact boundaries of masked regions (Fig. S18C) or choose random positions to mask in an allowed region on each iteration of inpainting. Ensembles can be generated using $RF_{joint2}$ , either in a manner akin to that described above, or by inpainting sequence/structure between arbitrary specified coordinates. For the example shown in Figure 5D, the coordinates of two residues were randomly translated up to a specified distance from their original positions, and the network was tasked with inpainting the masked region given the unmasked positions of the two translated residues.

### Supplementary Text

##### Structure prediction

The hallucination pipeline uses published versions of trRosetta (*11*) and RoseTTAFold (*15*), as well as a development version of RoseTTAFold with 2 tracks (MSA and pair features) rather than 3 tracks. For most of the designs shown, we used an improved variant of RoseTTAFold where the MSA transformer (*55*) track is replaced with a Perceiver (*56*).

Because RoseTTAFold only predicts backbone coordinates, we added sidechains to the output of a hallucination run using Rosetta and refined the full-atom structure by relaxing once in torsion space with predicted pairwise restraints and once in cartesian space with only pairwise distance restraints and C$\alpha$ coordinate restraints. Outputs from the trRosetta-based hallucination pipeline were relaxed similarly, except a structural model was first built by minimizing against the predicted pairwise restraints because trRosetta does not directly predict 3D coordinates. An optional minimization of RoseTTAFold backbone models against pairwise restraints prior to sidechain modeling was sometimes also performed for a small increase in structure quality. The output of the final relax step is the model used for downstream analysis and further design.

##### Coordinate RMSD loss

In addition to the cross-entropy motif loss, sometimes we used an additional RMSD motif loss $\mathcal{L}_{M,RMSD}$ defined as the backbone (N, C$\alpha$, C) root-mean-squared distance between predicted and reference motif coordinates after superposition(*57*). While using $\mathcal{L}_{M,RMSD}$ alone did not yield as good designs as using the cross-entropy loss $\mathcal{L}_M$ alone, a combination of the two losses (with weights $w_M = 1$ and $w_{M,RMSD} = 0.5$) gave the best DAN-lDDT and motif RMSD.

**KL divergence loss**

In some cases we defined the hallucination loss as a Kullback-Leibler (KL) divergence rather than entropy, following previous practice(*12*). Given network predictions $p(y)$ and background distributions $q(y)$ discretized over *B* bins,

$$\mathcal{L}_{H,KL} = -\sum_{y\in\{d,\omega,\theta,\phi,\theta^T\phi^T\}} \left[ \left( \sum_{i=1}^{L} \sum_{j\neq i}^{L} (1 - m_{ij}) \sum_{b=1}^{B} p(y_{ijk}) \log \frac{p(y_{ijb})}{q(y_{ijb})} \right) / \left( \sum_{i=1}^{L} \sum_{j\neq i}^{L} (1 - m_{ij}) \right) \right]$$

The background distributions represent residue-residue distance and angle distributions conditioned on only sequence separation, without knowledge of the amino acid identities(*11*, *58*). We generated the background using a separately trained neural network (for trRosetta) or by averaging the predictions for 100 random sequences (RoseTTAFold). The KL hallucination loss gave generally similar results as the entropy loss, although entropy yielded designs with higher helical content.

**Auxiliary losses**

For some problems we used additional auxiliary loss terms consisting of repulsive, attractive, and radius-of-gyration terms (Fig. S2):

$$\mathcal{L}_{aux} = w_{rep}\mathcal{L}_{rep} + w_{atr}\mathcal{L}_{atr} + w_{rog}\mathcal{L}_{rog}.$$

The repulsive and attractive losses $\mathcal{L}_{rep}$ and $\mathcal{L}_{atr}$ are partial Lennard-Jones potentials with a user-specified characteristic distance $\sigma$ (Fig. S2B)(*59*). The potentials are a function of the distance between predicted backbone atoms of the hallucinated protein and all atoms of a user-defined binding partner, and averaged over all such pairs (Fig. S2A).

The radius of gyration loss $\mathcal{L}_{rog}$ is used to control the overall shape of generated proteins and to indirectly favor a well-packed core (Fig. S2B). It is defined as an exponential linear unit with a user-specified threshold $R_0$:

$$\mathcal{L}_{rog} = \begin{cases} R_g, & R_g > R_0 \\ \exp(R_g - R_0), & R_g \leq R_0 \end{cases}$$

where the radius of gyration $R_g$ is calculated as the root-mean-squared position of the predicted C$\alpha$ positions $\boldsymbol{r}_{C\alpha}$:

$$R_g = \sqrt{\frac{1}{L}\sum_{i}^{L} \|r_{C\alpha}\|^2}$$

For epitope presentation and receptor decoy hallucinations, we used the repulsive and radius-of-gyration losses, with $w_{rep} = 1$, $\sigma_{rep} = 4$ Å, $w_{rog} = 1$, and $R_0 = 18$ Å. For binder design, we used both repulsive and attractive losses, with $w_{rep} = 1$, $\sigma_{rep} = 4$ Å, $w_{atr} = 10$, $\sigma_{atr} = 6$ Å,$w_{rog} = 1$, and $R_0 = 18$ Å. The unweighted attractive loss is typically 50-100x smaller than the other loss terms, so it is given a higher weight.

**Fixing and avoiding specific amino acids**

When residues on the functional motif are known to form desirable interactions with the binding partner or a ligand, we constrained these positions to stay the same (native) amino acid during optimization. Conversely, we also included the ability to avoid certain amino acids at all positions (e.g. cysteine). Both capabilities are implemented as adding or subtracting a large number ($10^8$) to the sequence logits at the beginning of each optimization.

When hallucinating di-iron proteins, we fixed the native amino acid identities of metal-coordinating residues buried in the core of a helical bundle (Fig. 3A, Table S1). This led to the serendipitous finding that hallucination can generate buried hydrogen bond networks (Fig. S10). (Fig. S10), likely because RoseTTAFold does not model the metal ligand and therefore tries to satisfy the buried polar groups with hallucinated polar residues. Although RoseTTAFold does not explicitly model sidechains, the sequences thus generated could be predicted by AF to have surprisingly dense hydrogen bond networks. These are difficult to design with traditional methods (*60*), and so hallucination could provide a novel solution to this problem.

**Automatic motif placement by exhaustive triplet enumeration**

The fixed motif-placement method described in the main methods is simple, but requires extensive sampling to identify good motif placements and iterative rounds of design to efficiently explore the search space. To avoid this sampling and iteration, we developed a method to automatically place the motif during optimization by means of a modified motif loss that rewards recapitulation of the motif in any location on the protein.

Consider a motif consisting of M discontinuous segments or “contigs” being placed on a protein of length L. Exhaustive enumeration of all contig placements in all possible positions in the designed sequence would require $O(L^M)$ loss evaluations and is not feasible for multi-segment motifs with many contigs (large M>3). However, the M=3 case is still practically realizable, so we developed an approach which exhaustively enumerates placements for all possible contig triplets from the motif and forces placements of different triplets to be self-consistent (described below). This was achieved by developing a two-term loss function

$$\mathcal{L}_{motif} = \mathcal{L}_{sat} + \mathcal{L}_{con}.$$

The first term $\mathcal{L}_{sat}$ forces recapitulation of the entire motif by averaging over $\binom{M}{3}$ $\mathcal{L}_{sat}^{abc}$ scores controlling how well each individual $abc$ triplet fits into the hallucinated structure:

$$\mathcal{L}_{sat} = 1/\binom{M}{3} \sum_{abc\in\{\text{triplets}\}} \mathcal{L}_{sat}^{abc}$$

Given network predictions $p(y)$, triplet-wise satisfaction scores $\mathcal{L}_{sat}^{abc}$ are calculated as a weighted average of cross entropy scores $H_{ijk}^{abc} = H_{ij}^{ab} + H_{jk}^{bc} + H_{ik}^{ac}$ for placing contigs a,b,c at positions i,j,k in the sequence:

$$\mathcal{L}_{sat}^{abc} = \sum_{i,j,k} p_{ijk}^{abc} H_{ijk}^{abc}$$

where $H_{ij}^{ab} = -\sum_{y\in\{d,\omega,\theta,\phi,\theta^T,\phi^T\}} \log p(y_{ij} = y^0)$, $y_{ij}$ is the predicted distance or orientation angle between positions i and j, and $y^0$ is the desired value of the geometric parameter between contigs a and b. The best placement of 3 contigs is the (i,j,k) that minimizes $H_{ijk}^{abc}$. To favor emergence of a single best placement during optimization, we weight the 3-body cross entropy scores by their statistical weight:

$$p_{ijk}^{abc} = exp\,(-\beta H_{ijk}^{abc}) / \sum_{i,j,k} \exp(-\beta H_{ijk}^{abc})$$

The inverse temperature parameter $\beta$ controls the strength of constraints and is increased throughout optimization from 2 to 20.

In the triplet decomposition, it is possible that different triplets $abc$ and $abd$ sharing a pair of contigs $ab$ may yield optimal placements $ijk$ and $lmn$ such that$ij \neq lm$. To discourage this, we use a “triplet consistency” loss defined as the negative symmetrized cross-entropy between marginal probabilities of placements of a given contig pair in different triplets, averaged over all order 4 permutations of contigs $a, b, c, d$:

$$\mathcal{L}_{con} = -1/\binom{M}{4} \sum_{a,b,c,d\in\{quadruplets\}} \frac{1}{L^2} \sum_{i,j} \left( p_{ij}^{ab(c)}\, log\, p_{ij}^{ab(d)} + p_{ij}^{ab(d)}\, log\, p_{ij}^{ab(c)} \right)$$

where $p_{ij}^{ab(c)} = \sum_k p_{ijk}^{abc}$ is the probability of placing 2 contigs a,b at positions i,j marginalized over the placements of a 3rd contig c.

During optimization, we use $\mathcal{L}_{sat} + \mathcal{L}_{con}$ instead of $\mathcal{L}_M$ as the motif loss term, and at the end of optimization we identify contig placements by looking for high-scoring cliques in the weighted adjacency matrix $A_{ij} = p_{ij}^{ab(c)} + p_{ij}^{ac(b)} + p_{ij}^{bc(a)}$ averaged over all triplets a,b,c.

**Automatic motif placement by greedy search**

Although fully differentiable, the triplet enumeration method above requires $O(L^3M^3)$ memory for a length L protein with M contigs, and only approximately computes the motif loss when $M > 3$. An alternative method uses a greedy search algorithm to place the contigs and applies the exact motif loss given this placement. On each optimization step, this method will:

1. Place 2 of the contigs by computing the cross-entropy between their inter-contig geometries and network predictions at all possible starting residue numbers i and j

where they can be placed, respectively. Try this for all pairs of contigs and keep the placement of the 2 contigs with the lowest score.

2. Place remaining contigs one at a time, minimizing its inter-contig cross entropy with the already-placed contigs, until all have been placed.

3. Once all contigs are placed, the standard motif loss is calculated for that placement and used to compute the gradient.

Contigs are required to remain in a user-defined order. Positions that would result in the contigs overlapping with each other or prevent the placement of the remaining contigs are scored as positive infinity. Because greedy searches can miss global optima, we added the top 3 scoring results at each step to a search tree, yielding a collection of possible contig placements.

Intuitively, this method will initially place contigs in positions that randomly happen to score well, but after a few gradient updates, these regions will match the contigs more and more and the process becomes self-reinforcing. Because this method only evaluates pairs of contigs, it requires $O(L^2)$ memory and $O(M)$ time.

##### RF*joint* training loss formulation

The loss function formulation for RF*joint* is as follows. Note that structure related losses are applied over the entire predicted protein, and the sequence cross entropy loss is only applied at masked regions.

$$\mathcal{L}_{total} = 0.05\mathcal{L}_{dist} + 1\mathcal{L}_{aa} + 0.025\mathcal{L}_{tors} + 0.5\mathcal{L}_{FAPE} + 0.05\mathcal{L}_{bond\ angle} + 0.05\mathcal{L}_{bond\ length} + 0.05\mathcal{L}_{lddt}$$

Where $\mathcal{L}_{dist}$ is a cross entropy loss over the distogram and anglegram as described in (*15*), predictions $\mathcal{L}_{aa}$ is a cross entropy loss over any masked positions in the input MSA, $\mathcal{L}_{tors}$ is a cross entropy loss on binned backbone dihedral angle predictions, $\mathcal{L}_{FAPE}$ is a backbone level frame aligned point error, as described in (*16*), with a relu cutoff of 20. $\mathcal{L}_{bond\ angle}$ is calculated as $\mathcal{L}_{bond\ angle} = \sum_{i=1}^{L}\left(\sqrt{\left(\hat{\theta}_{Ca_i,C_i,N_{i+1}} - \theta_{Ca_i,C_i,N_{i+1}}\right)^2} + \sqrt{\left(\hat{\theta}_{C_i,N_{i+1},Ca_{i+1}} - \theta_{C_i,N_{i+1},Ca_{i+1}}\right)^2}\right)$ where $\hat{\theta}_{a_i,b_j,c_k}$ is the planar angle between atoms *a*, *b*, and *c* from residues *i*, *j*, and *k* (respectively) resulting from a backbone prediction, $\theta_{a_i,b_j,c_k}$ is the ideal planar bond angle between those atoms, and *L* is the number of amino acids in the protein. $\mathcal{L}_{bond\ length}$ is calculated as $\mathcal{L}_{bond\ length} = \sum_{i=1}^{L}\sqrt{(\hat{D}_{C_i,N_{i+1}} - D_{C_i,N_i+1})^2}$ where $\hat{D}_{a_i,b_j}$ and $D_{a_i,b_j}$ are the predicted and ideal covalent bond lengths between atoms *a* and *b* from residues *i*, and *j*, respectively. $\mathcal{L}_{lddt}$ is the lDDT loss as calculated in (*15*).

##### PD-1 mimetic design and yeast display

We used trRosetta to hallucinate 100,611 designs scaffolding a 2-segment beta-sheet motif from the HAC PD-1 interface (5IUS chain A residues 63-82, 119-140), and selected those with

DAN-lDDT > 0.6 and interface RMSD < 1.5 Å, or 66,501 designs. These were sequence-designed using the Rosetta FastDesign mover with layer design and fragment based PSSMs to constrain amino acid choices in the protein. We also constrained interface residues (chain A 64, 66, 68, 70, 73-75, 77-78, 81, 85, 89-91, 124, 126, 128, 132, 134, 136, 139) to only repack (keeping only native amino acids at these positions) and put harmonic coordinate restraints to these residues to ensure they didn't move during relaxation. We then filtered designs on a panel of Rosetta- and deep-learning-based metrics capturing quality of the binder monomer, the sequence-structure consistency, and the quality of the interface (Fig. S1A), selecting 3,042 designs for experimental testing (*4*, *13*, *61*). Designs were reverse-translated and split in 2 halves to be synthesized by Twist on a 300-bp oligonucleotide chip, assembled by PCR, cloned into pETCON3 for Aga2p and c-Myc fusion, and transformed into yeast strain EBY100 for surface display (*4*).

For yeast surface display, the pool of transformed yeast was inoculated into CTUG medium (yeast nitrogen base 6.7g/L (difco) + complete amino acids -trp -ura + 2% glucose) and incubated 12-16 hours at 30°C with shaking, then diluted 200uL + 2mL into SGCAA (yeast nitrogen base 6.7g/L + complete amino acids 5g/L (Bacto) + 90mM $Na_2HPO_4$ + 2% galactose + 0.1% glucose) and incubated 12-16 hours to induce binder expression and display. For flow sorting, around $10^7$ cells were harvested, washed 3x in TBSF (50mM Tris-HCl pH8.0, 150mM NaCl, 1% bovine serum albumin), incubated in TBSF with biotinylated PD-L1 (R&D Systems) for 30 minutes at room temperature, washed 1x in TBSF, incubated for 30 minutes at room temperature in 0.1mg/mL FITC anti-c-Myc (ICL Lab) and 70mg/mL streptavidin R-phycoerythrin (PE) conjugate (Invitrogen), and washed 3x in TBSF. The PD-L1 and FITC/PE were added in the same incubation when labeling with avidity. Cells were sorted on a Sony SH800 flow sorter and $10^4$ - $10^6$ FITC+/PE+ cells were collected (Fig. S1B), and cultured in CTUG again for the next sort. A series of 4 sorts were performed: first twice at 1uM PD-L1 with avidity, then 1uM, and then at 100nM, 10nM, and 1nM (Fig. S1B). Cells from the final sort were plated on CTUG agar plates and 56 colonies were Sanger sequenced to identify the designs (Fig. S1C).

The 3 most abundant designs were validated by labeling clonal yeast cultures in a titration of PD-L1 and measuring on an Attune NxT flow analyzer (Invitrogen). The resulting data was processed by manually choosing a FITC threshold for expression (log10 FITC > 3.2) and fitting a hyperbola $y = A\frac{x}{x+K}$ with free parameters *A, K* to the mean PE-H/FITC-H, where *A* is the maximal binding signal and *K* is the apparent $K_d$. Plots in (Fig. S1E) are shown with data normalized to fitted *A* so all curves saturate at 1. Competition experiments with unlabeled wildtype PD-1 were performed with clonal yeast cultures of the binders in a similar manner (Fig. S1F).

**Native protein scaffold search**

How necessary was it to use deep learning methods to scaffold the chosen functional sites? Without the ability to generate protein backbones, we would be limited to grafting the motifs into existing native protein backbones. To estimate the difficulty of using this non-deep learning approach, we used the Rosetta MotifGraft (*62*) mover to search the PDB for proteins that we could potentially graft the motifs into, with the following requirements:

1. The resulting chimera does not clash with the binding target (if applicable).
2. For motifs with multiple segments, all of the starting and ending backbone atoms of the motif have a combined RMSD less than 1A to the graft sites in the native proteins. For added flexibility, each motif segment could replace a native segment up to 50 aa longer or shorter than itself. (ie - A short loop in the motif could replace a long native loop, so long as the starting and ending points were close to each other.)
3. For motifs with only a single element, the corresponding native segment be the same length and have an all backbone RMSD less than 1A. We could not filter on just the RMSD of the starting and ending points because it poorly constrains the orientation of the amino acids, resulting in non-plausible chimera junctions.

To account for sequence (and structural) redundancy, the sequences of all single protein chains in the Protein Data Bank (PDB) solved by x-ray crystallography were clustered at a 30% sequence identity threshold using mmseqs2 (*63*) and assigned to a unique cluster. The number of suitable native scaffolds reported in Table S3 is the number of clusters that had at least one match, excluding the cluster that the motif was taken from. The frequency is that number divided by the total number of clusters in the PDB.

The number of matches is still likely an overestimate, since many of the matched native proteins are highly structurally homologous to the original structure the motif was taken from and therefore unlikely to scaffold the motif in a meaningfully different way. Additionally, there is no constraint that the matched native proteins be small or compact, an advantage in potentially downstream applications and a requirement that the hallucinated designs generally fulfill.

For the RSV-F site V motif, we performed a more detailed analysis against both the PDB100 and the AlphaFold proteomes database (*34*) (Fig. S13). The motif, which contains a single contiguous segment, was searched against all possible positions in all structures (or models) in the 2 databases, and the lowest backbone RMSD was recorded for each structure. We filtered out any structure containing clashes (defined as heavy atoms closer than 2Å) to the antibodies against this motif in PDB:5TPN, as well as any whose sequence was more than 50% identical to 5TPN. Only 2 results (6w16, 5wb0) remained with RMSDs lower than our best designs (or a frequency of $2/(355712+161370) = 3.9\times10^{-6}$ across the 2 databases), and even these are distantly related (36% identity) to 5TPN and highly related (90% identity) to each other. Before filtering out homologs and receptor clashes, we obtained 67 scaffolds in the databases (frequency $1\times10^{-4}$) better than our best design.

##### Using RFjoint to design helical repeat proteins

As an additional test of the utility of our inpainting approach, we used it to generate designed helical repeat proteins (DHRs), a widely useful class of proteins requiring diverse interfaces with precise periodicity (*64*). To design novel DHRs, we generated an ensemble of $RF_{Joint}$ inpaintings. Symmetry was achieved by providing a small tri-helix fragment from an existing DHR (5CWJ, 5CWH). Inpaintings with high AF2 prediction lDDT and RMSD to the model were then fused into a repeat protein.

##### AlphaFold hallucination for benchmarking

To generate the AF hallucinations used for the benchmarking analysis in Fig. S21, we used AF model 5 (model_5_ptm) to perform 3000-5000 steps of MCMC to minimize the mean predicted alignment error TM-score as a loss, starting from a random starting sequence with an initial acceptance temperature of 0.08 and halflife of 500 steps, mutating randomly first 3 (step 1-800), then 2 (step 801-1600), then 1 (step 1601-end) amino acids at a time, the highest TM-score model across the entire trajectory was chosen as final model. These are purely hallucinated proteins with no functional site, analogous to (*12*).

### Supplementary Figures

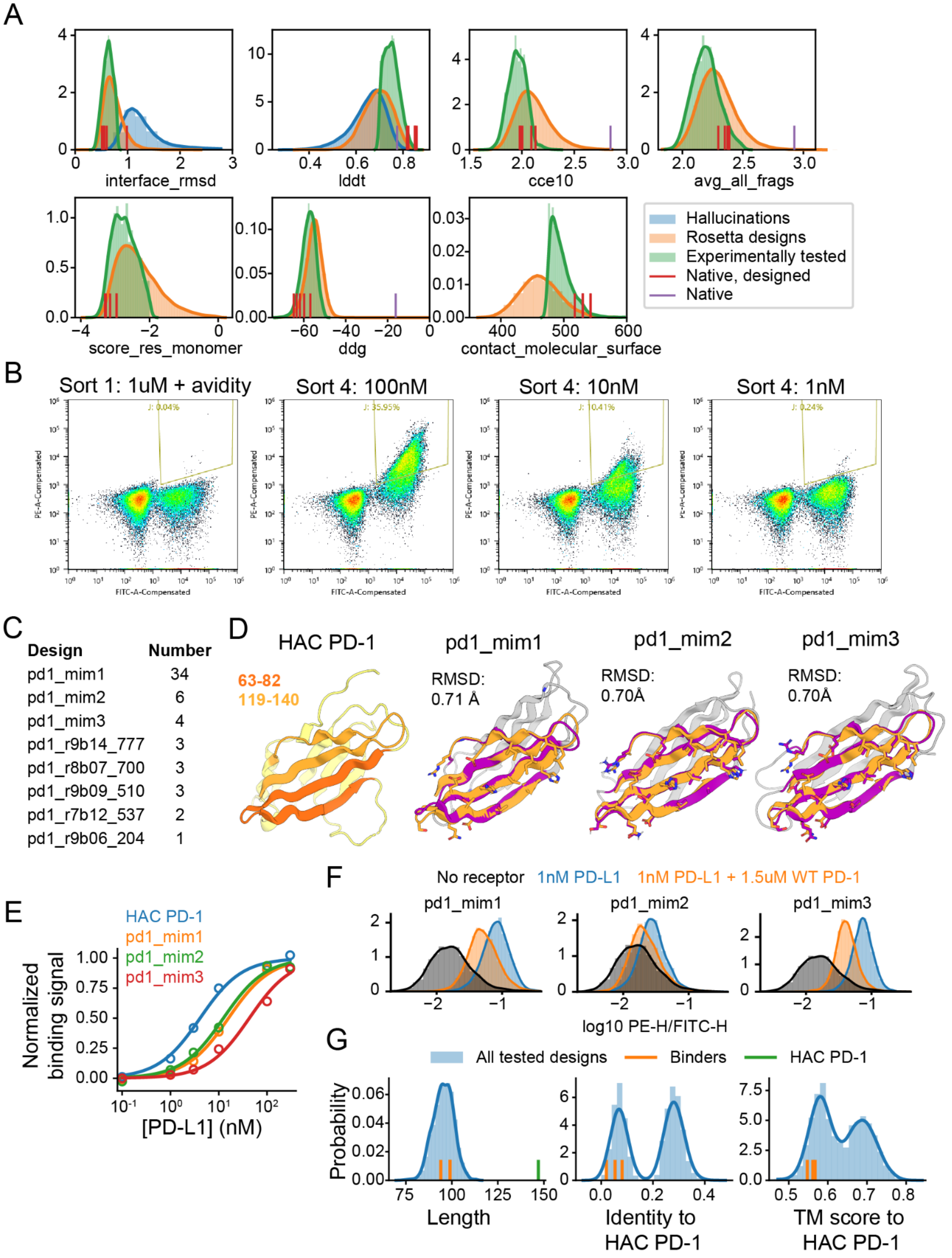

**Figure S1. Design and testing of PD-1 mimetics**

(A) Distributions of metrics for PD-1 hallucinations, Rosetta designs, and experimental library. Interface_rmsd is C$\alpha$ RMSD over 22 interface positions (Supplementary Text), lddt is DeepAccNet predicted lDDT, cce10 is cross-entropy of residue-residue distances and angles of the design model to trRosetta predictions for the design sequence, filtered to pairs within 10 Å (*13*), avg_all_frags is a measure of local sequence-structure agreement (*13*), score_res_monomer is the Rosetta energy per residue for the hallucinated binder, ddg is a Rosetta-estimated $\Delta\Delta G$ of binding, and contact_molecular_surface is a measure of the interface area (*4*). We selected designs for testing which had: interface RMSD < 0.8 Å and (DAN-lDDT > 0.75 or cce10 < 2.1 or avg_all_frags < 2.1) and DAN-lDDT > 0.7 and avg_all_frags < 2.5 and score_res_monomer < -2 and contact_molecular_surface > 475 and ddg < -50. (B) PE (binding) vs FITC (surface displayed protein) signal during FACS sorting of PD-1 mimetics. Sort 2 (1uM PD-L1 with avidity) and 3 (1uM PD-L1, no avidity) are not shown. (C) Number of colonies matching the sequence of a given design among 56 colonies that were Sanger sequenced. (D) Crystal structure of HAC PD-1 (discontinuous interface motif in 2 shades of orange) and design models of 3 experimentally isolated binders. “RMSD” denotes the C$\alpha$ RMSD between design model and template motif at 22 interface residues (Methods). (E) Binding signal (Methods) from clonal yeast cultures versus receptor concentration for HAC PD-1 and designs isolated from pooled sorting. Apparent $K_d$ values in nM are: HAC PD-1: 4.10; pd1_mim1: 15.9; pd1_mim2: 12.5; pd1_mim3: 42.9. (F) Normalized PE (binding) signal for clonal yeast cultures expressing the 3 binders in the presence of receptor and receptor + unlabeled purified wildtype PD-1. (G) Distribution of sequence length, amino-acid identity to HAC PD-1, and TM-score to HAC PD-1 for the 3,038 experimentally tested designs. The values for HAC PD-1 and the 3 binders shown in (D) are plotted as vertical bars.

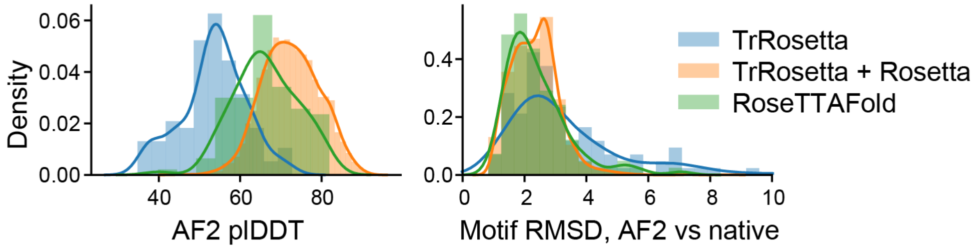


**Figure S2. Comparison of trRosetta and RoseTTAFold for hallucinating PD-1 mimetics**

AlphaFold predicted lDDT and motif backbone RMSD (AF model versus native motif) for hallucinations generated using trRosetta, RoseTTAFold, or trRosetta followed by Rosetta-based sequence design.

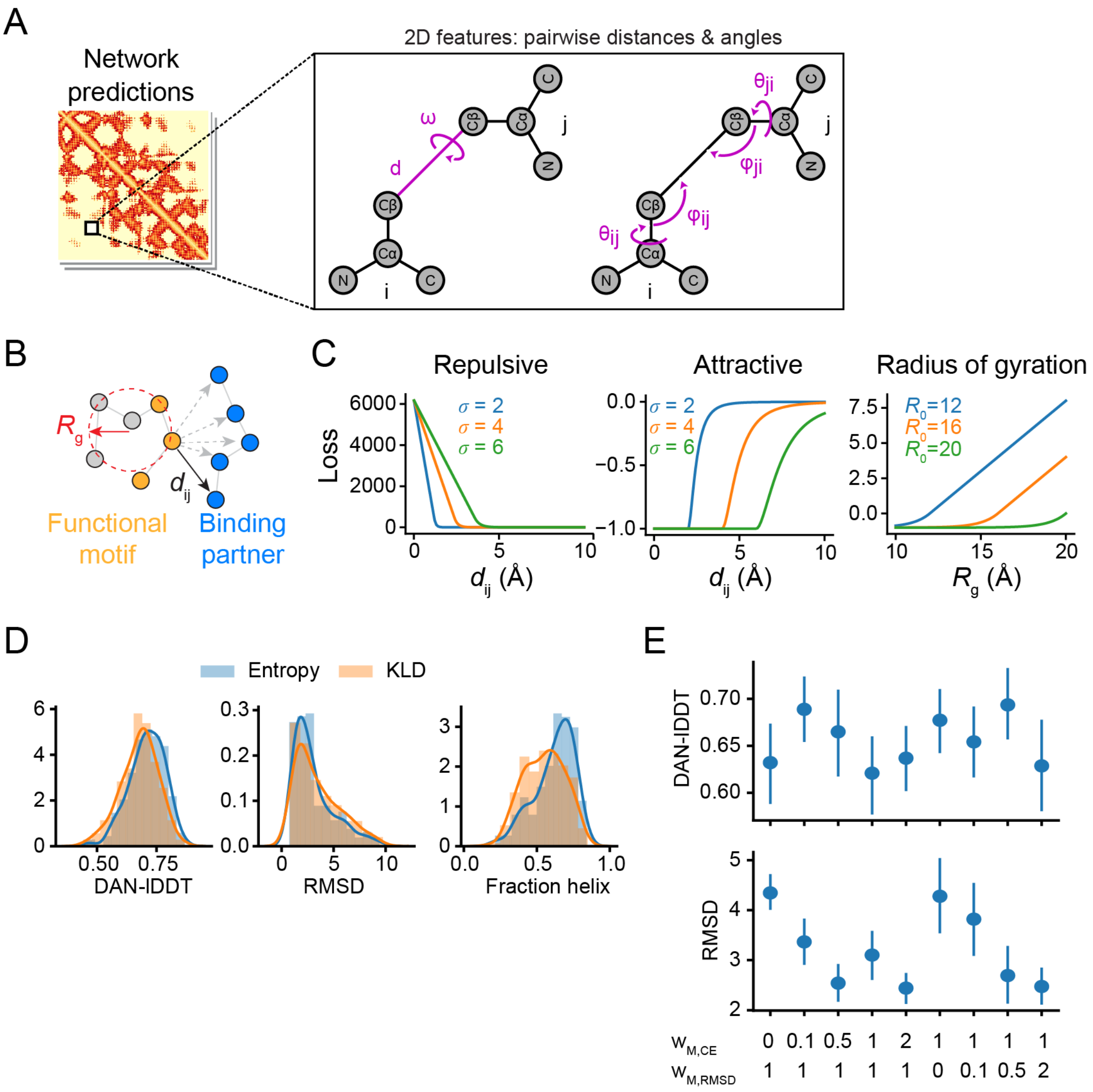


**Figure S3. Auxiliary and alternative loss terms**

(A) Schematic of the pairwise distances and orientation angles whose distributions are predicted by trRosetta and RoseTTAFold and which are used to define the motif and hallucination losses. (B) Schematic of radius of gyration and distances used to calculate repulsive and attractive losses (Supplementary Text). (C) Functional forms of the losses. (D) Distributions of DAN-lDDT, motif RMSD, and fraction of residues that are helix for designs generated using entropy or KL divergence hallucination losses (Supplementary Text), for scaffolding a 2-segment motif from C3d (1GHQ chain A residues 104-126, 170-185). (E) DAN-lDDT and motif RMSD for the same C3d scaffolding problem as in (D), but with varying the loss term weights for the cross-entropy based motif loss ($w_{M,CE}$) or RMSD-based motif loss ($w_{M,RMSD}$).

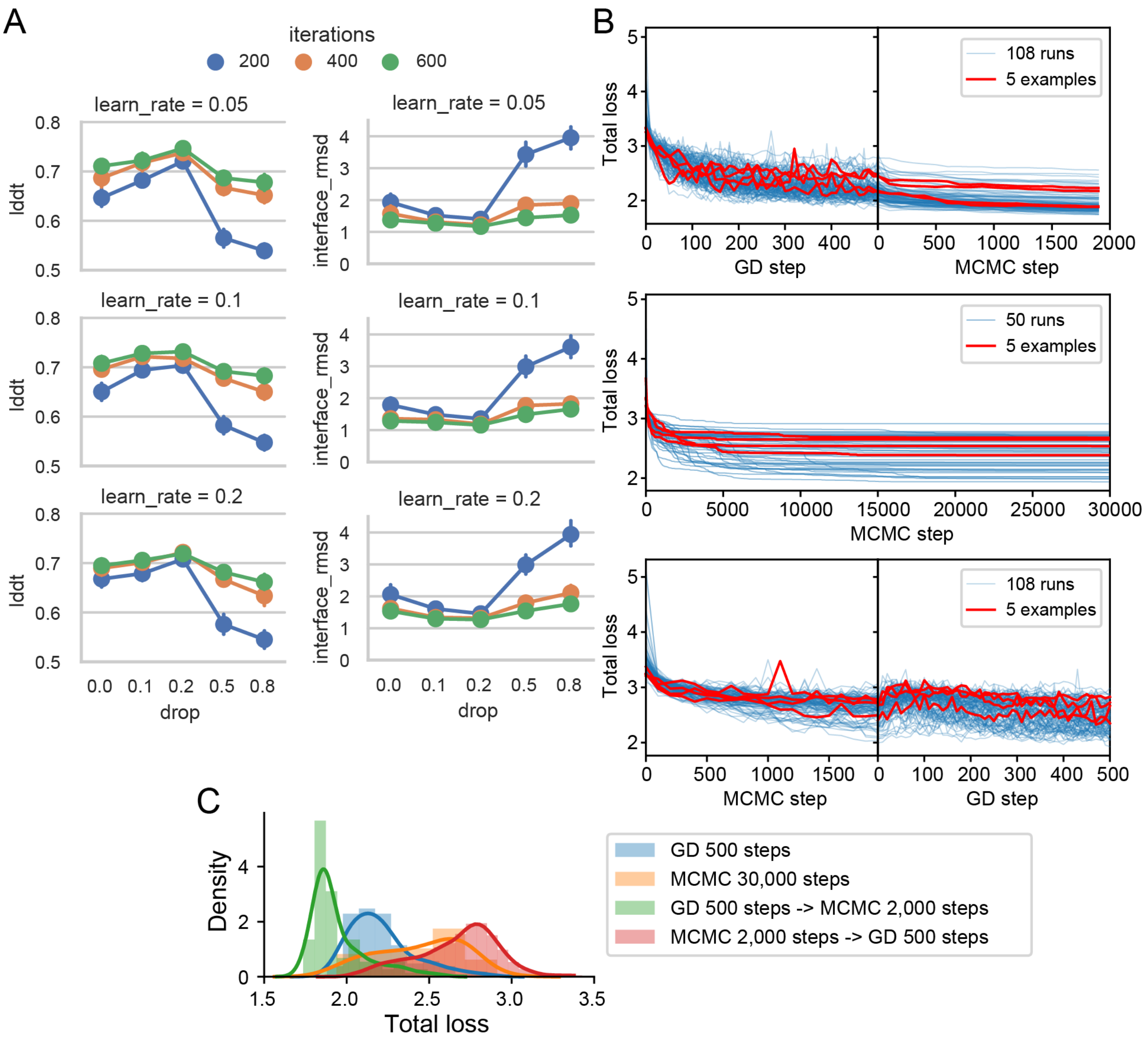


**Figure S4. Optimization by gradient descent and MCMC**

(A) DAN-lDDT and C$\alpha$ RMSD at 22 interface residues for a hyperparameter scan for gradient descent using RoseTTAFold with a 2-segment motif from HAC PD-1 (residues 63-82 and 119-140). Plotted are mean and 90% confidence interval of 50-100 trajectories per condition. Optimal parameters were dropout = 0.2, learning rate = 0.05. Running 600 iterations gave the best results but 400 steps was comparable and therefore used for most problems. (B) Loss trajectories for gradient descent (GD) followed by MCMC, MCMC only, or MCMC followed by GD. (C) Distributions of final losses for the trajectories shown in (B).

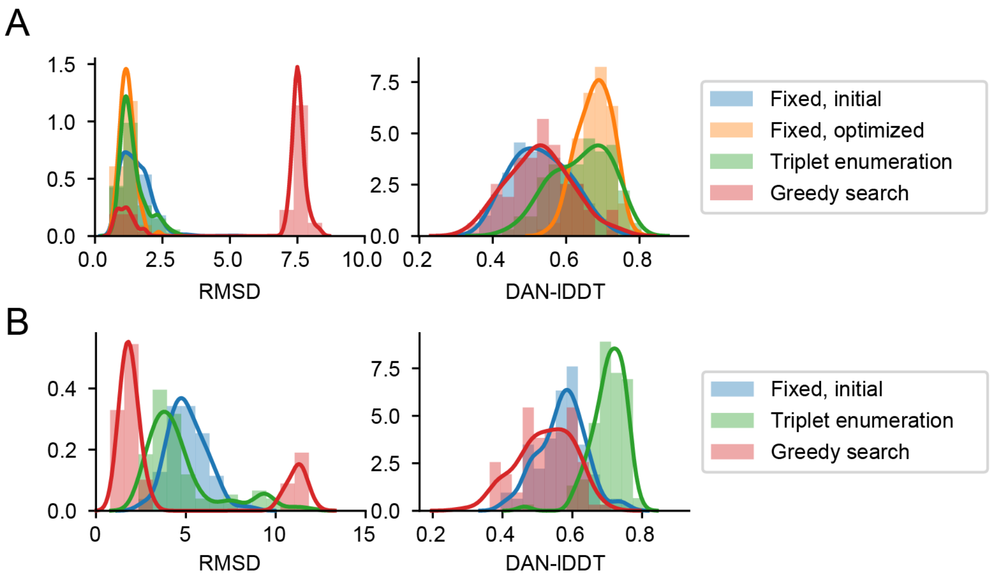


**Figure S5. Comparison of motif-placement methods**

Distributions of Motif C$\alpha$ RMSD and DeepAccNet-predicted lDDT for (A) PD-1 mimetics hallucinated with different motif placement methods (Methods, Supplementary Text). Motif consists of 2 discontinuous segments as shown in Fig. 2. “Fixed, initial” is an initial run of the fixed motif placement method where contigs are placed anywhere along a given length, and “Fixed, optimized” is a run where gaps between contigs are chosen based on results from the “initial” run. (B) Same methods but with a 2-segment motif from C3d (1GHQ: A104-126,A170-185) and a “fixed, optimized” run was not done.

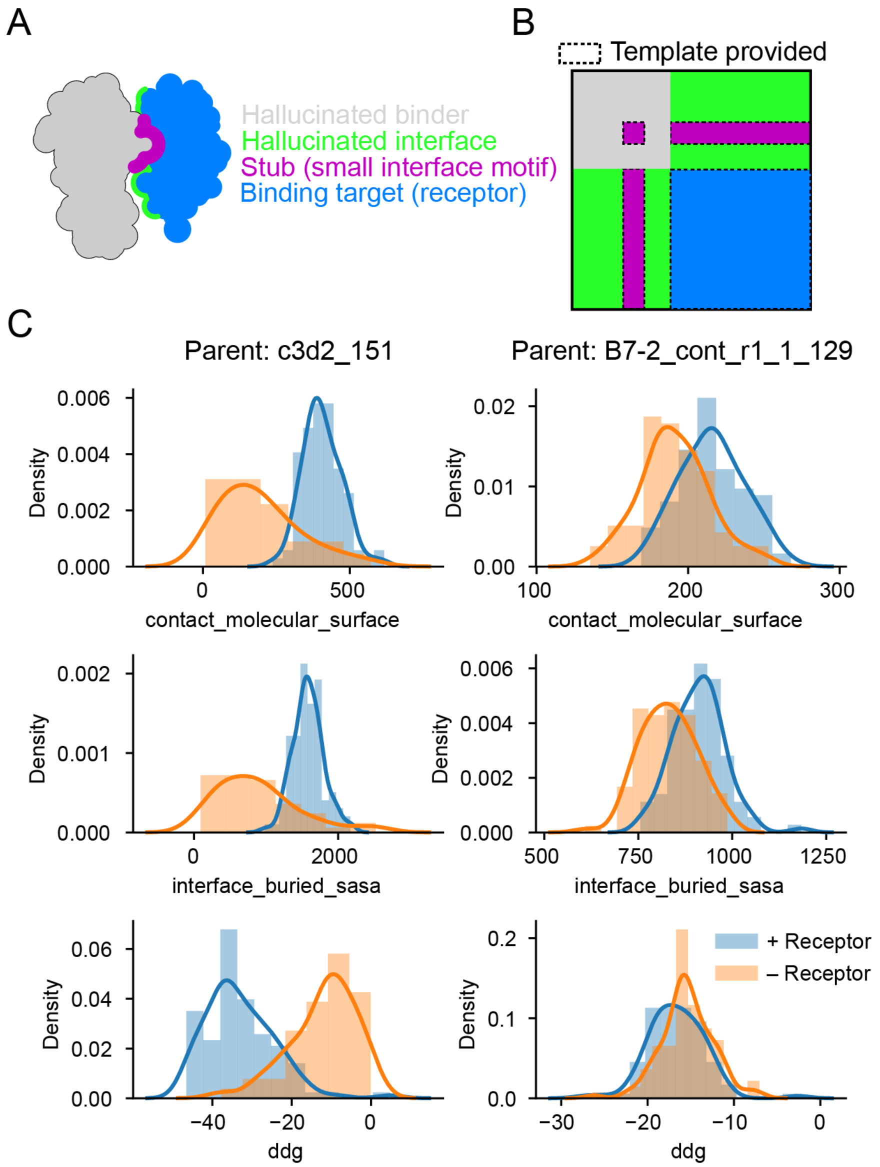


**Figure S6. Multi-chain hallucination for binder refinement**

(A) Schematic of the different regions of a hallucinated binder and its binding partner. (B) Pairwise features from RoseTTAFold that are considered during two-chain interface refinement. A template input is given for the entire binding target, the pre-defined (native) stub, and their relative geometry. (C) Interface metrics for refined B7-2 and C3d mimetics, with or without explicit modeling of the binding partner.

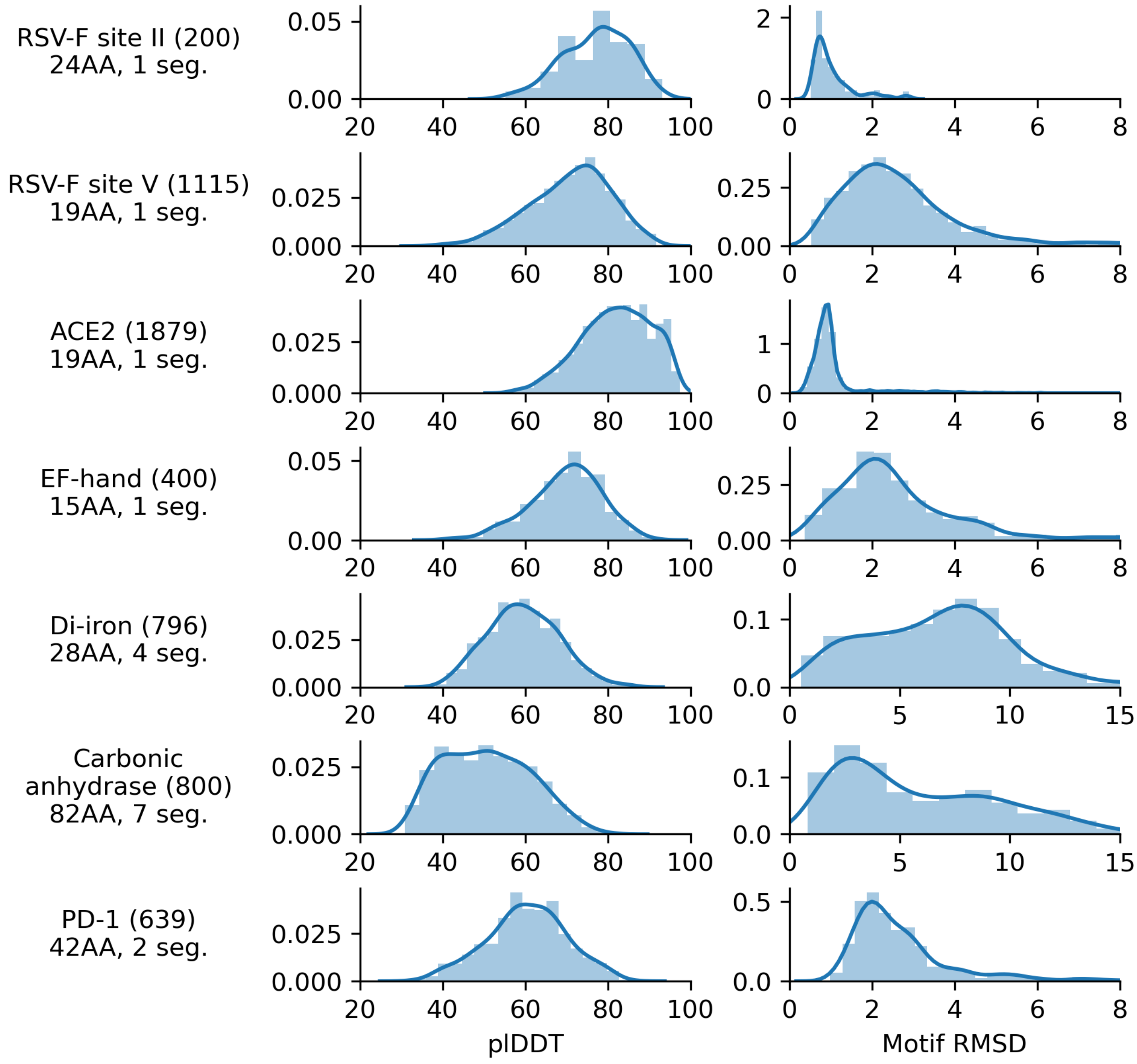


**Figure S7. Distribution of pIDDT and motif RMSD of hallucinations before filtering**

Distributions over all hallucinated (single-chain) designs of (A) AlphaFold pIDDT and (B) backbone RMSD between native motif and AF predictions from hallucinated sequences. Numbers in parentheses indicate the number of designs being plotted.

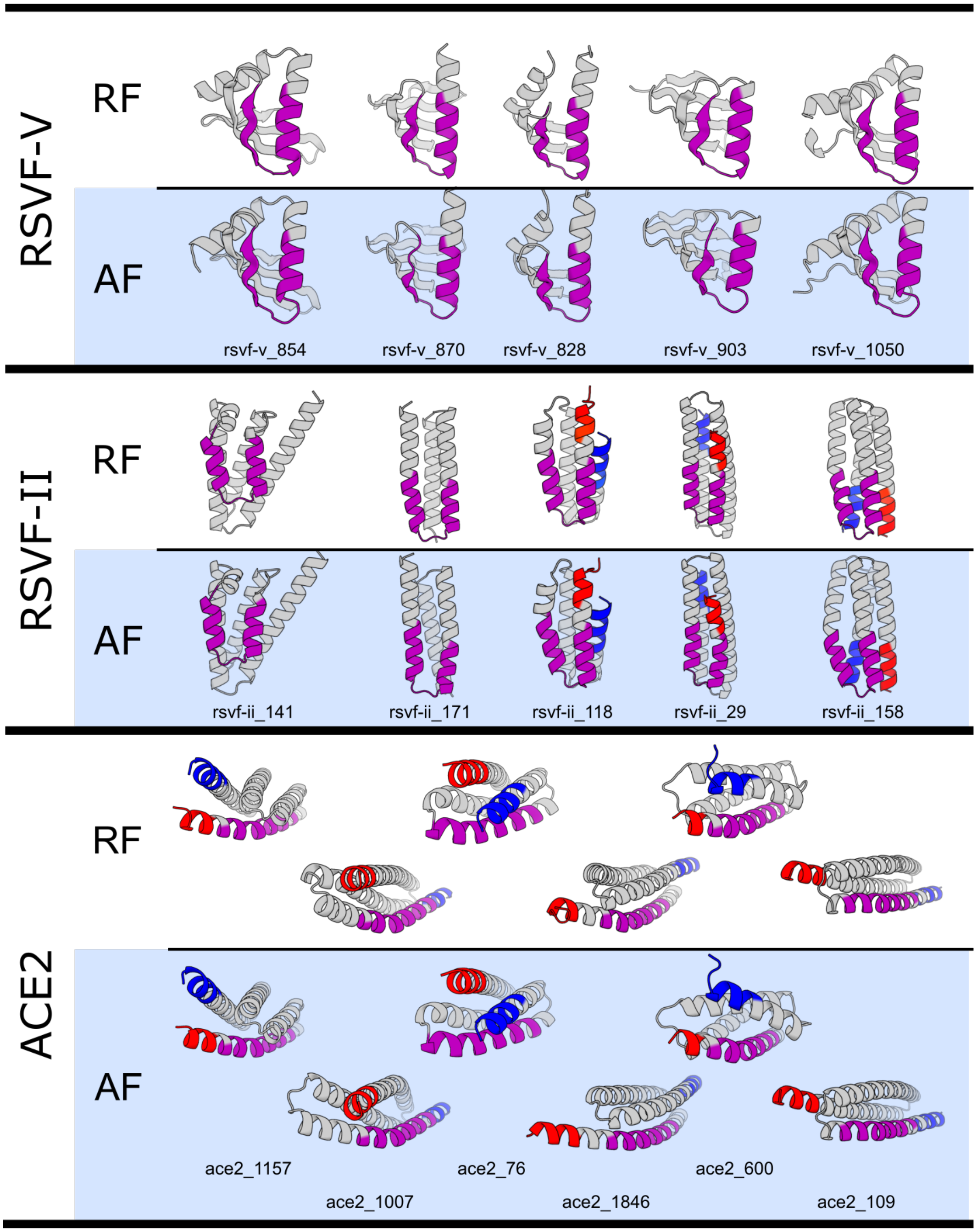


**Figure S8. RoseTTAFold and AlphaFold models of epitope scaffolds and receptor traps**

Main text and additional RosettaFold (RF) hallucinations and their corresponding AlphaFold (AF) model are shown for epitope scaffolds and receptor trap design problems. Functional motifs are highlighted in purple. The N- and C-termini in some designs have been colored blue and red (respectively) to highlight that hallucination can find diverse topological solutions, despite having similar overall folds. Detailed metrics for these designs can be found in Table S2.

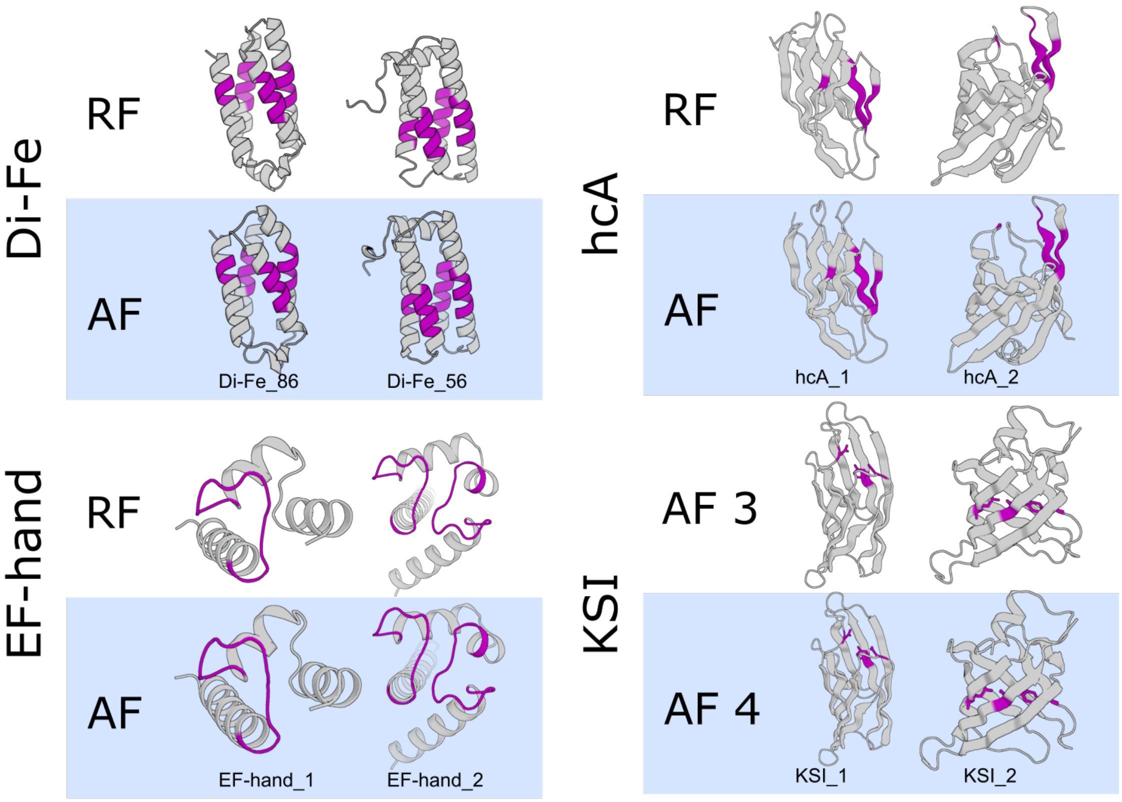


**Figure S9. RoseTTAFold and AlphaFold models of metal-binding and active sites**

Main text active site hallucinations and their corresponding AlphaFold (AF) model are shown. Functional motifs are highlighted in purple. Detailed metrics for these designs can be found in Table S2.

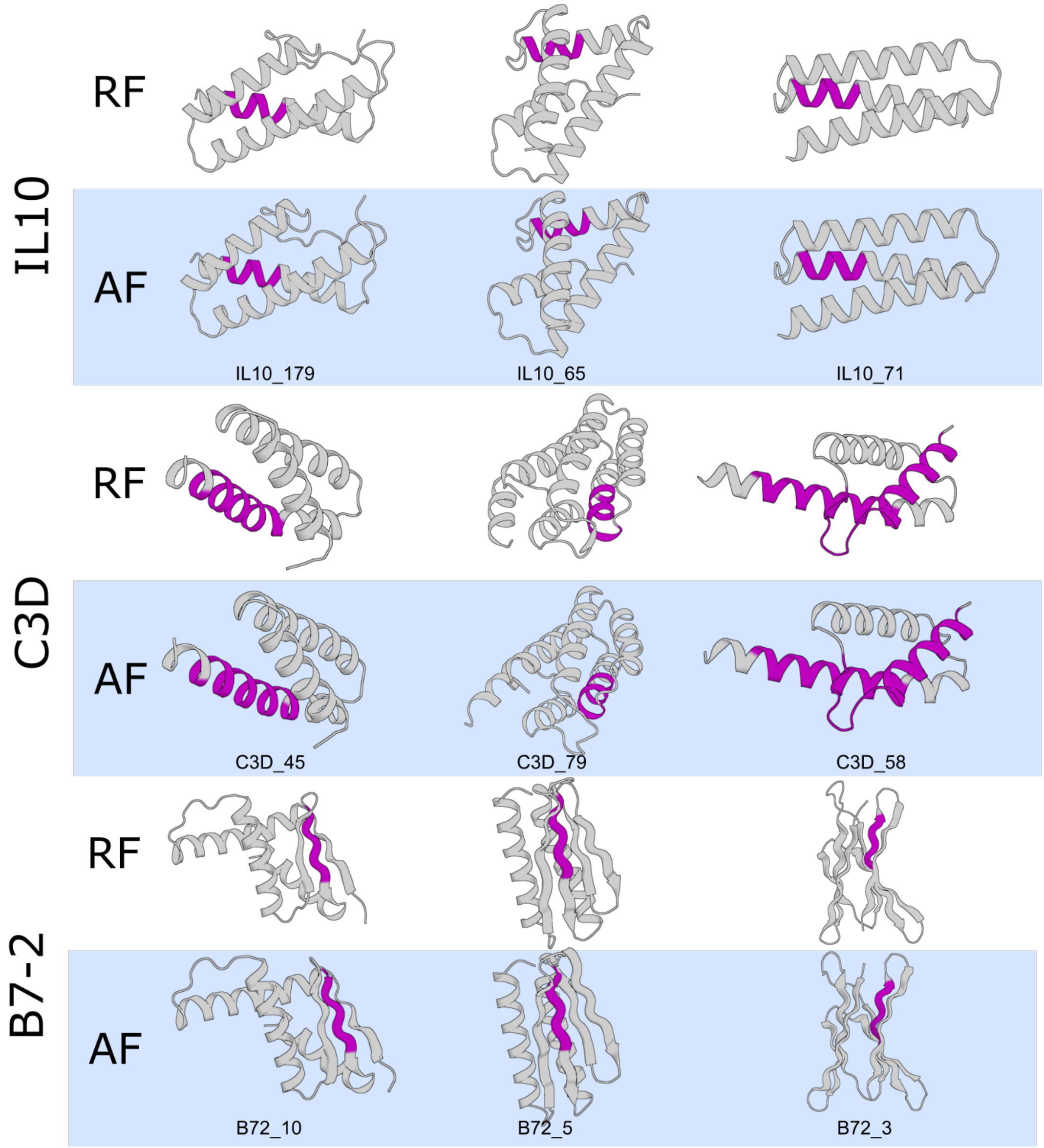


**Figure S10. RoseTTAFold and AlphaFold models of PPI designs**

Main text PPI hallucinations and their corresponding AlphaFold (AF) model are shown. Functional motifs are highlighted in purple. Detailed metrics for these designs can be found in Table S2.

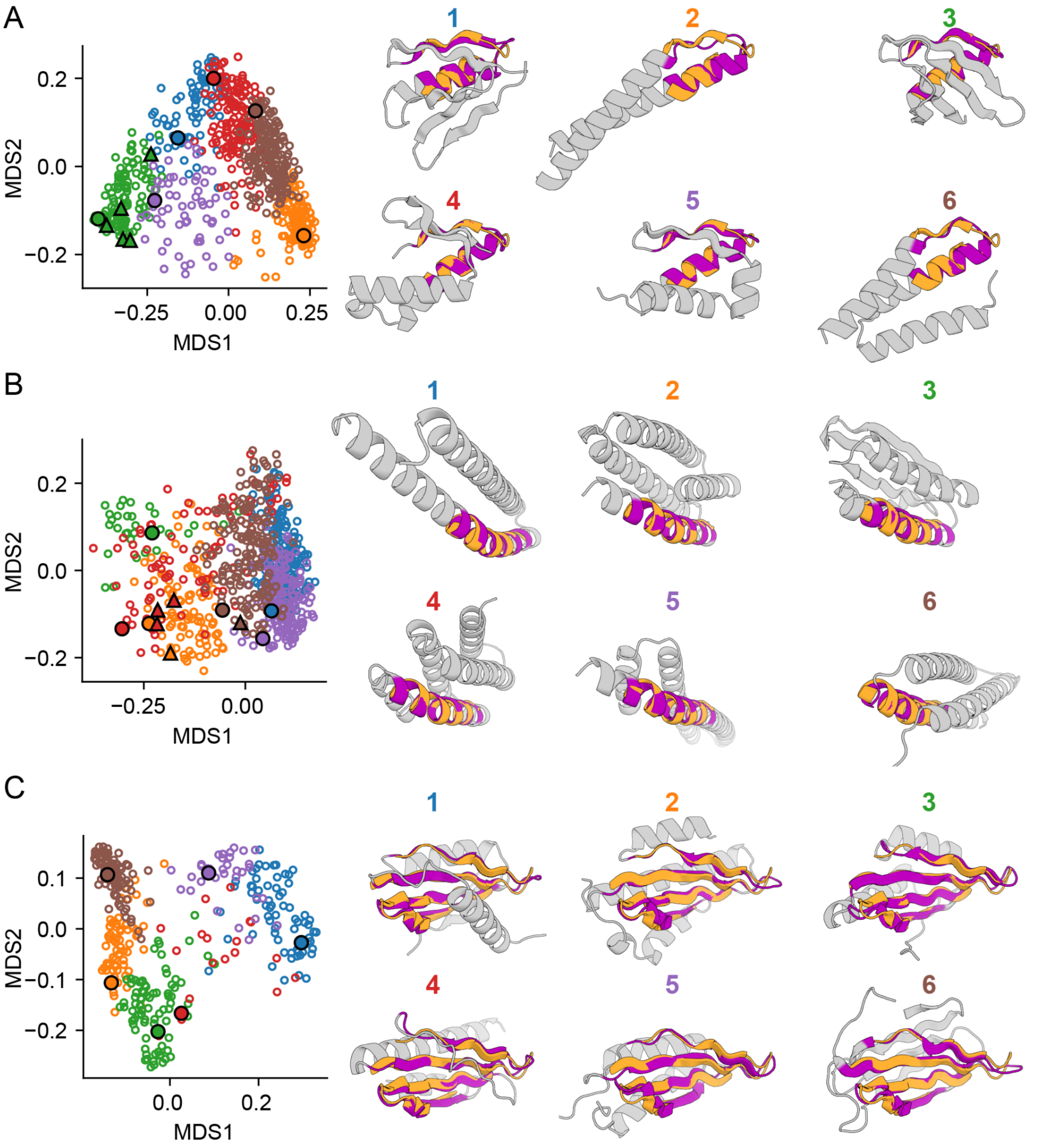


**Figure S11. Structural diversity of hallucinated designs**

Structural diversity of hallucinated (A) RSV-F site V scaffolds, (B) ACE2 mimics, and (C) PD-1 mimics. Raw hallucinations were filtered for AF pIDDT > 60 and motif RMSD (AF vs native) < 6 Å (removing 25%-50% of designs) and all pairwise TM-scores were computed and projected into 2 dimensions using classic multidimensional scaling (left scatterplots). 6 clusters were identified using k-means, and cluster representatives (black-outlined circles) were selected manually and shown on the right. The number of clusters was chosen arbitrarily. Triangles represent designs shown in Fig. 2-3 and S8. Structures are colored to highlight native motif (orange) and motif in hallucination (purple).

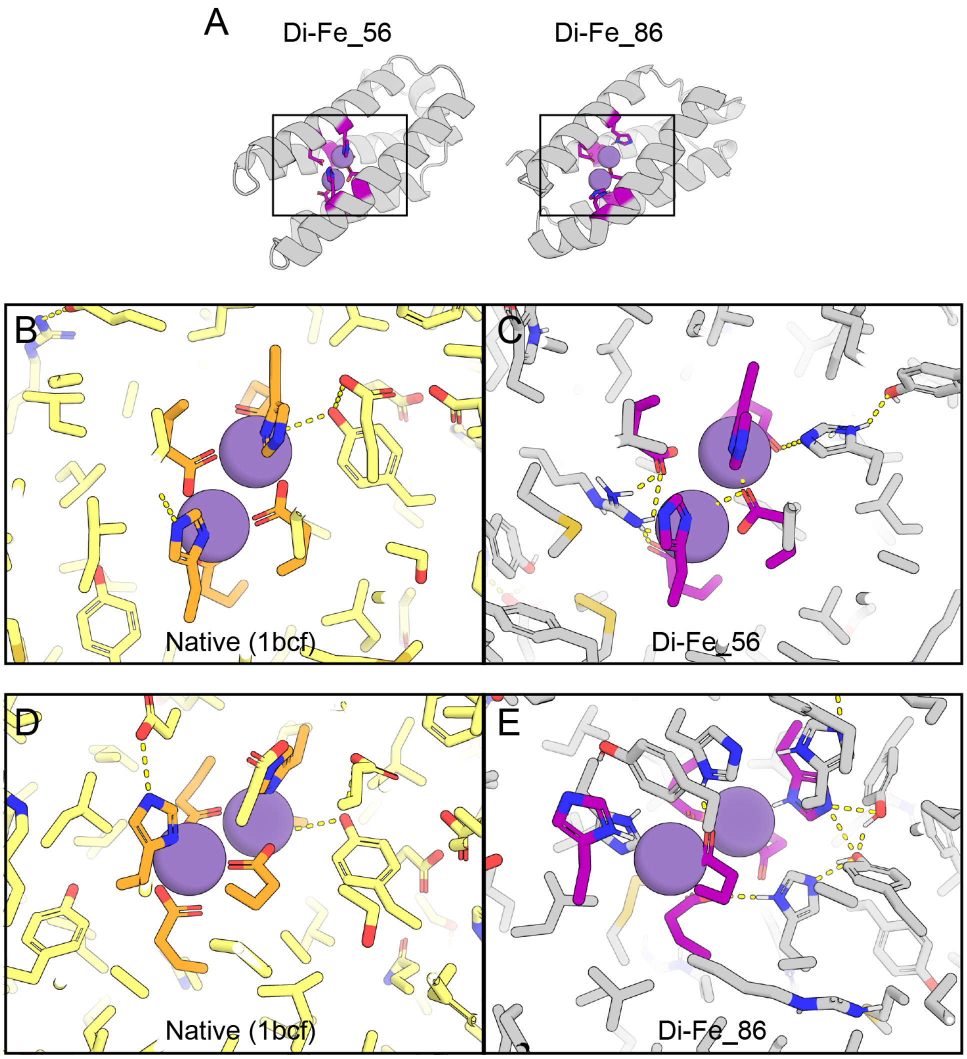


**Figure S12. Hallucinations containing buried hydrogen-bond networks**

(A) Two di-iron hallucinations (also shown in Fig. 3) and close-ups (C, E) of the residues near the metal binding site. Structures are AF predictions after AMBER relax (*65*). The native protein used as a hallucination reference is shown in (B, D) after aligning to the hallucinations on the backbone atoms of the functional residues (orange in native, purple in hallucinations). Metals shown in (C, E) are taken from the native structure after superimposition. Note the presence of hallucinated polar residues (gray histidines and tyrosines) to form hydrogen-bonding networks with the functional histidines and glutamates, which were constrained to their native identities during hallucination.

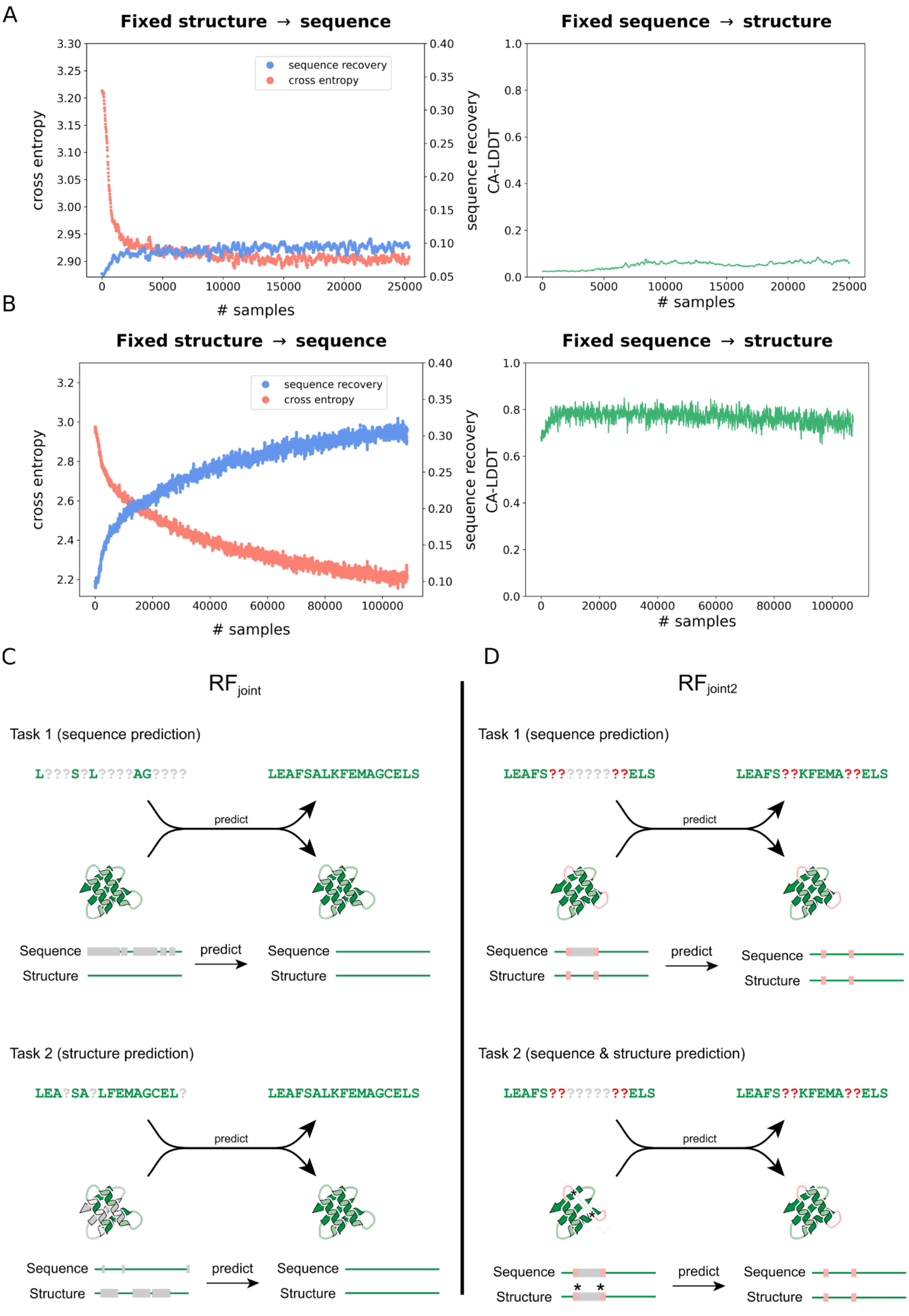

A
Fixed structure → sequence
sequence recovery
cross entropy
cross entropy
sequence recovery
### samples
Fixed sequence → structure
CA-LDDT
### samples
B
Fixed structure → sequence
sequence recovery
cross entropy
cross entropy
sequence recovery
### samples
Fixed sequence → structure
CA-LDDT
### samples
C
RFjoint
Task 1 (sequence prediction)
L???S?L????AG????
LEAFSALKFEMAGCELS
predict
Sequence
Structure
predict
Sequence
Structure
Task 2 (structure prediction)
LEA?SA?LFEMAGCEL?
LEAFSALKFEMAGCELS
predict
Sequence
Structure
predict
Sequence
Structure
D
RFjoint2
Task 1 (sequence prediction)
LEAFS?????????ELS
LEAFS??KFEMA??ELS
predict
Sequence
Structure
predict
Sequence
Structure
Task 2 (sequence & structure prediction)
LEAFS?????????ELS
LEAFS??KFEMA??ELS
predict
Sequence
Structure
predict
Sequence
Structure

**Figure S13. Training of joint sequence-structure recovery RoseTTAFold.**

Training curves showing the evolution of performance on fixed backbone sequence design and classic structure prediction while training Joint RosettaFold ($RF_{joint}$) using Algorithm S1. (A) Beginning from a completely untrained RosettaFold (RF) model, RF is unable to learn both sequence design and structure prediction tasks, as seen by saturation of the training curves at poor losses. Training was stopped early due to the saturation. (B) Starting from a RF model that has been pre-trained on only structure prediction, the model is able to learn sequence design well, while maintaining its structure prediction capabilities. This is accomplished using the same Algorithm S1. (C - D) Comparison of the training regimes for $RF_{joint}$ and $RF_{joint2}$, both of which were trained from a pretrained RF model. (C) $RF_{joint}$ was trained on two tasks. 75% of the time (Task 1), 90-100% of the sequence was masked, and the network tasked with recovering the sequence given the input template. 25% of the time (Task 2), the network was trained on the classic structure prediction task, with 15% of the MSA randomly masked, and some partial input templates provided. (D) $RF_{joint2}$ was trained on three tasks. 25% of the time (Task 1), the network was tasked with predicting the sequence of a contiguous masked region (10-25 residues long, see methods), given an input structure template, without the sequence or structure of 3-5 residues either side of the masked region. 50% of the time (Task 2), an identical contiguous 10-25 residue region was masked from the input sequence and template structure, plus 3-5 extra flanking residues on either side, and the network simultaneously tasked with predicting sequence and structure of the central (gray) masked region. The residue coordinates were provided for the residues either side of the masked region (see methods). The remaining 25% of the time, the network performed the structure prediction task illustrated in (C), bottom panel.

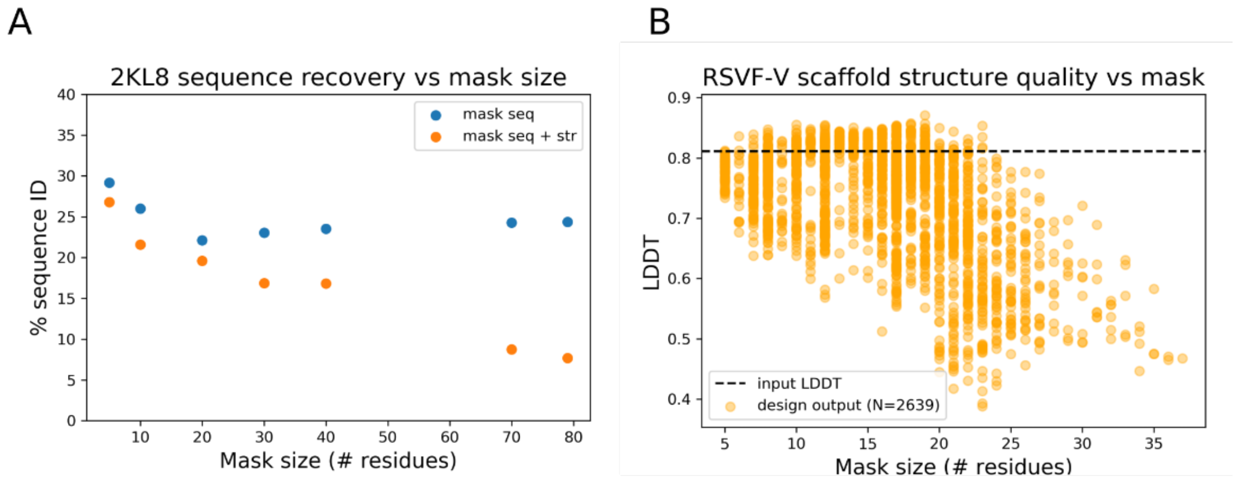


**Figure S14. Joint RosettaFold sequences and structures versus mask size**

Divergence of $RF_{joint}$ output sequences and structures from the input is tunable by changing the input mask size. (A) For test set *de novo* protein 2KL8, high sequence recovery to the native is achieved when masking small regions of sequence and structure. As the mask size is increased, sequence identity to the native decays. The limit of masking the entire sequence and structure is reached at 79 residues. (B) Given a hallucination output for the RSVF-V scaffolding problem, inpainting with $RF_{joint}$ can be used to diversify or idealize the protein. When masking very little sequence and structure, the output design DeepAccNet lDDT (*61*) is clustered very close to the input. In intermediate mask lengths (~5-22) output design LDDTs can be improved or deteriorated over the input design. In the very large mask regime (25+), DAN LDDTs monotonically decay, which is consistent with Figure S15.

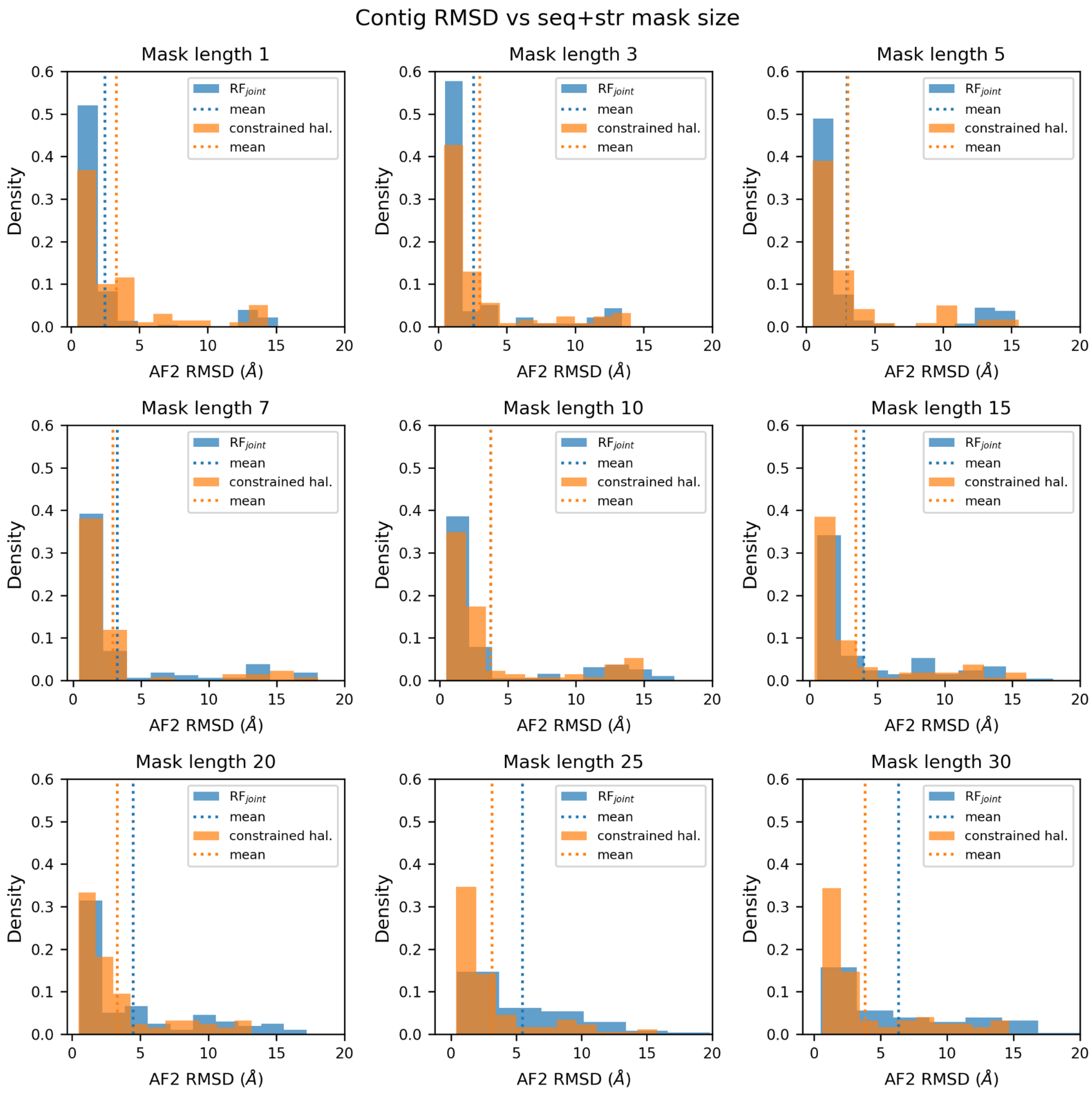


**Figure S15. Template RMSD vs starting information**

RMSD of a constrained region between a design and its AF2 prediction when designing a solution to a region missing both sequence and structure for a set of 10 *de novo* proteins, using either constrained hallucination (method 1) or RF$_{\text{joint}}$ missing information recovery (method 2). In regimes where more starting information is provided (i.e., mask lengths 1, 3, 5), information recovery with RF$_{\text{joint}}$ produces designs which better hold the input template structure in place, according to AF2. In the medium input information regime (i.e., mask lengths 7, 10, 15), RMSDs on the input templates are very similar between the two methods. In the low input information regime (i.e., mask lengths 20, 25, 30+), constrained hallucination outperforms RF$_{\text{joint}}$ information recovery.

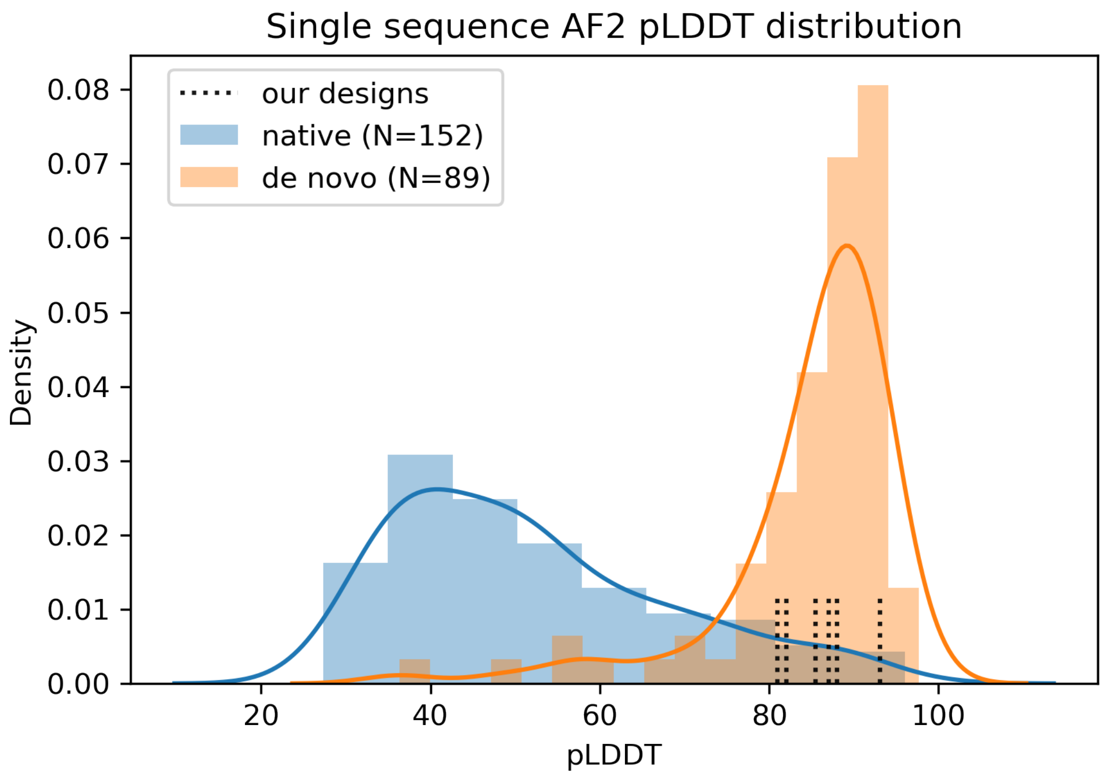


**Figure S16. Comparison of AF2 pLDDTs to other native or *de novo* proteins**

The strength with which a single amino acid sequence encodes its structure is a well known mechanism for filtering *de novo* protein designs down to a pool of high quality candidates for experimental testing (*4*, *13*). It is thus desirable for a protein design algorithm to generate sequences strongly encoding their structure. Here we compare the AF2 single sequence pLDDTs between a set of 152 native proteins (*66*), 89 structurally validated *de novo* proteins, and some of our designs from this study. We observe the designs obtained from our approaches match the distribution of high pLDDTs achieved by previous *de novo* proteins.

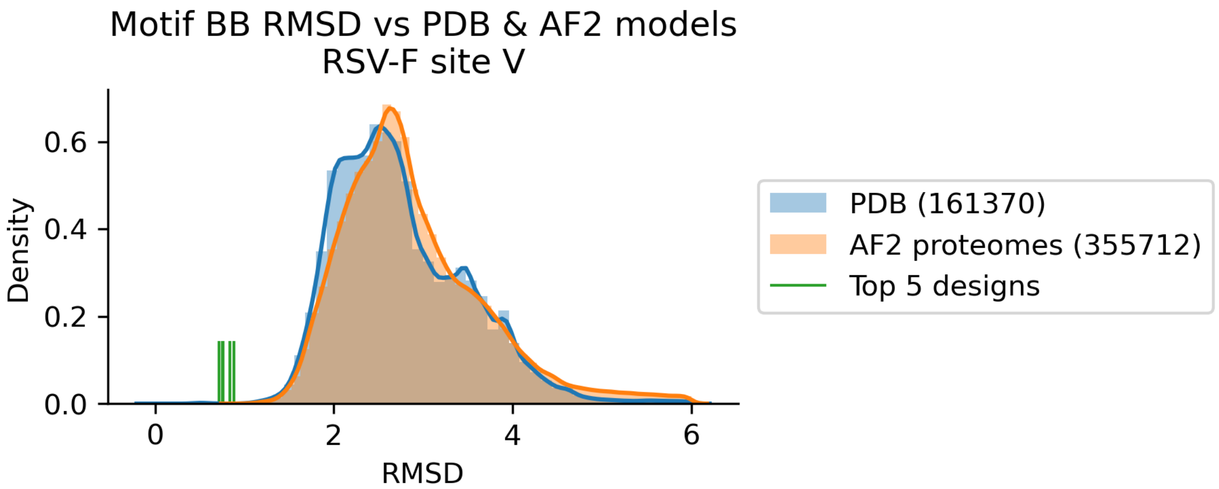


**Figure S17. RMSD of RSV-F motif hits in PDB and AlphaFold proteomes database**
Distribution of best RMSD between the RSV-F epitope (PDB 5tpn, chain A residues 163-181) and each structure in the PDB or AlphaFold2 proteomes database. The motif RMSD of the best 5 hallucinated designs are plotted for comparison. The frequency of finding an RMSD as good as any of these designs or better was $3.9 \times 10^{-6}$ (see "Native protein scaffold search" in Supplementary Text).

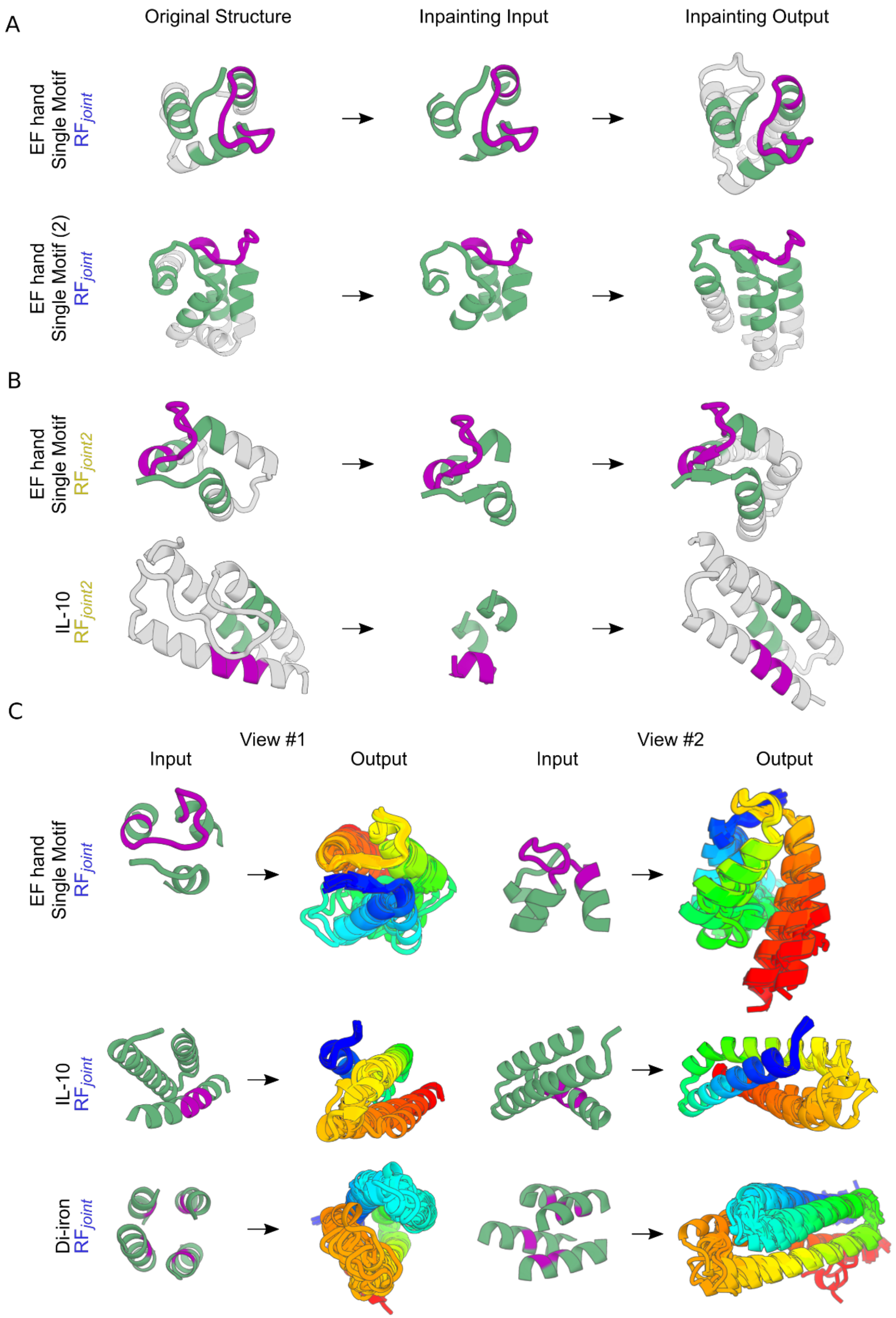
A
Original Structure
Inpainting Input
Inpainting Output
EF hand Single Motif RF joint
EF hand Single Motif (2) RF joint
B
EF hand Single Motif RF joint2
IL-10 RF joint2
C
View #1
View #2
Input
Output
Input
Output
EF hand Single Motif RF joint
IL-10 RF joint
Di-iron RF joint

**Figure S18. Additional $RF_{joint}$ / $RF_{joint2}$ designs and diversification**

Design of proteins harboring functional motifs via information recovery using $RF_{joint}$ and $RF_{joint2}$, and their diversification via changing the precise input masking pattern. All designs here are starting from a functional site (purple) embedded in a constrained hallucination (green if revealed to $RF_{joint}$ before design, gray if masked). (A) $RF_{joint}$ functional motif design examples. From top to bottom row with (AF2 motif RMSD to native, AF2 pLDDT): Single motif EF hand (87.5, 0.71), another single motif EF hand starting from a different hallucination (85.5, 0.71). (B) $RF_{joint2}$ functional motif design examples, starting from a smaller input template. Single motif EF hand (79.3, 0.88 Å), IL-10 (80.1, 0.44 Å). (C) Demonstration of the diversity of outputs obtained from an individual starting template via perturbing an input mask before designing with $RF_{joint}$. For an arbitrary mask with N contiguous segments, where the $i^{th}$ segment has an allowed variable length range of $A_i$ residues on its N-terminal end $B_i$ residues on its C-terminal end, the total number of unique masks one can create scales as $\prod_i A_i B_i$. This set of unique masks then provides means to perform a set of unique forward passes through $RF_{joint}$, and thus a set of unique designs. These designs can then be filtered downstream as desired.

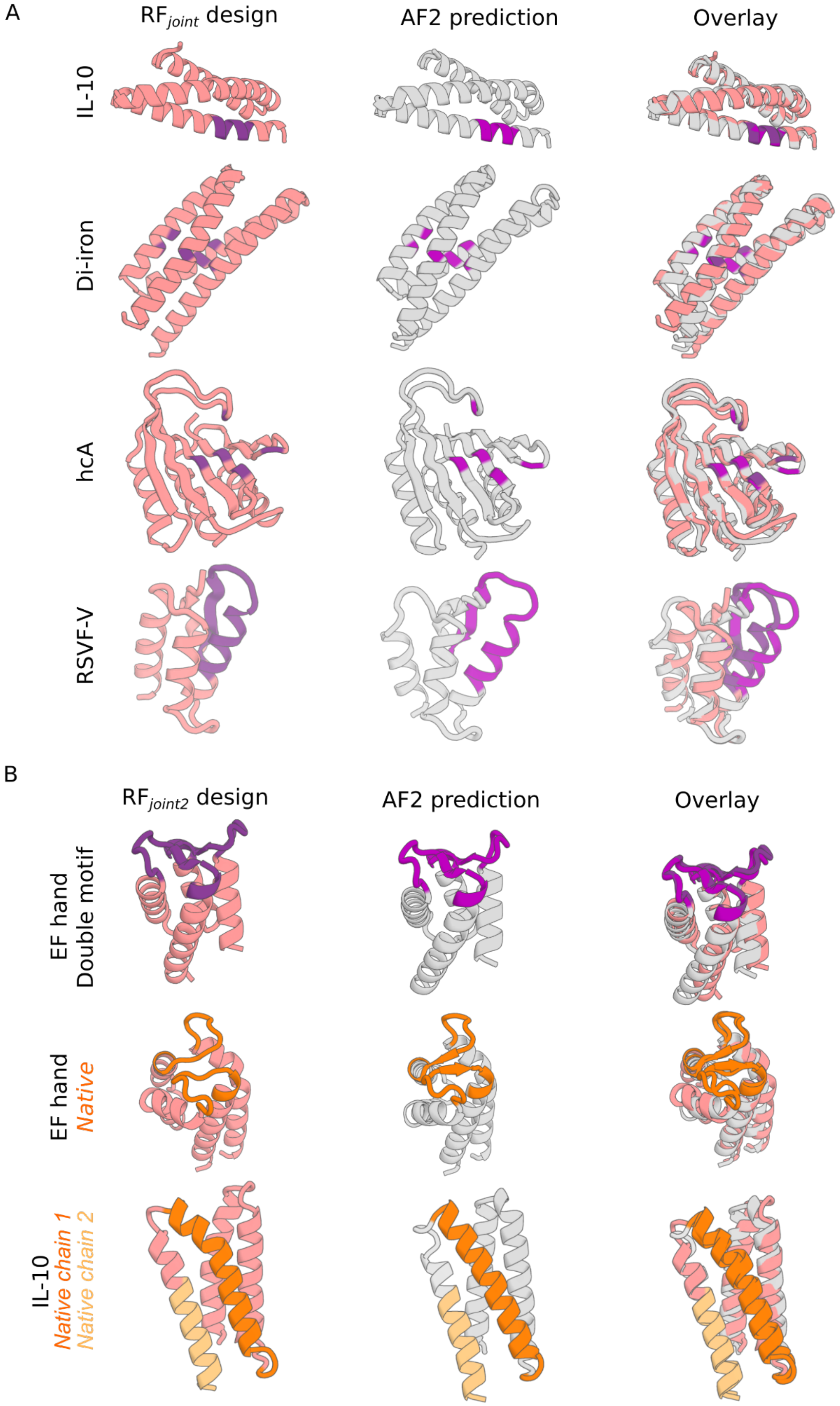
A
$RF_{joint}$ design
AF2 prediction
Overlay
IL-10
Di-iron
hcA
RSVF-V
B
$RF_{joint2}$ design
AF2 prediction
Overlay
EF hand
Double motif
EF hand
*Native*
IL-10
*Native chain 1*
*Native chain 2*

**Figure S19. Overlay of $RF_{joint}$ / $RF_{joint2}$ designs with AF2 predictions**

Overlays of the $RF_{joint}$ / $RF_{joint2}$ generated designs from Fig. 6, with structure predictions from AF2 (RMSD RF vs AF2). (A) $RF_{joint}$ inpainted designs scaffolding the IL-10 (1.86 Å), di-iron (1.04 Å), hcA (1.45 Å) and RSVF-V (2.76 Å) motifs. (B) $RF_{joint2}$ inpainted designs scaffolding the hallucinated EF hand double motif (0.82 Å), native EF hand double motif (1.92 Å) and the native IL-10 motif (1.47 Å).

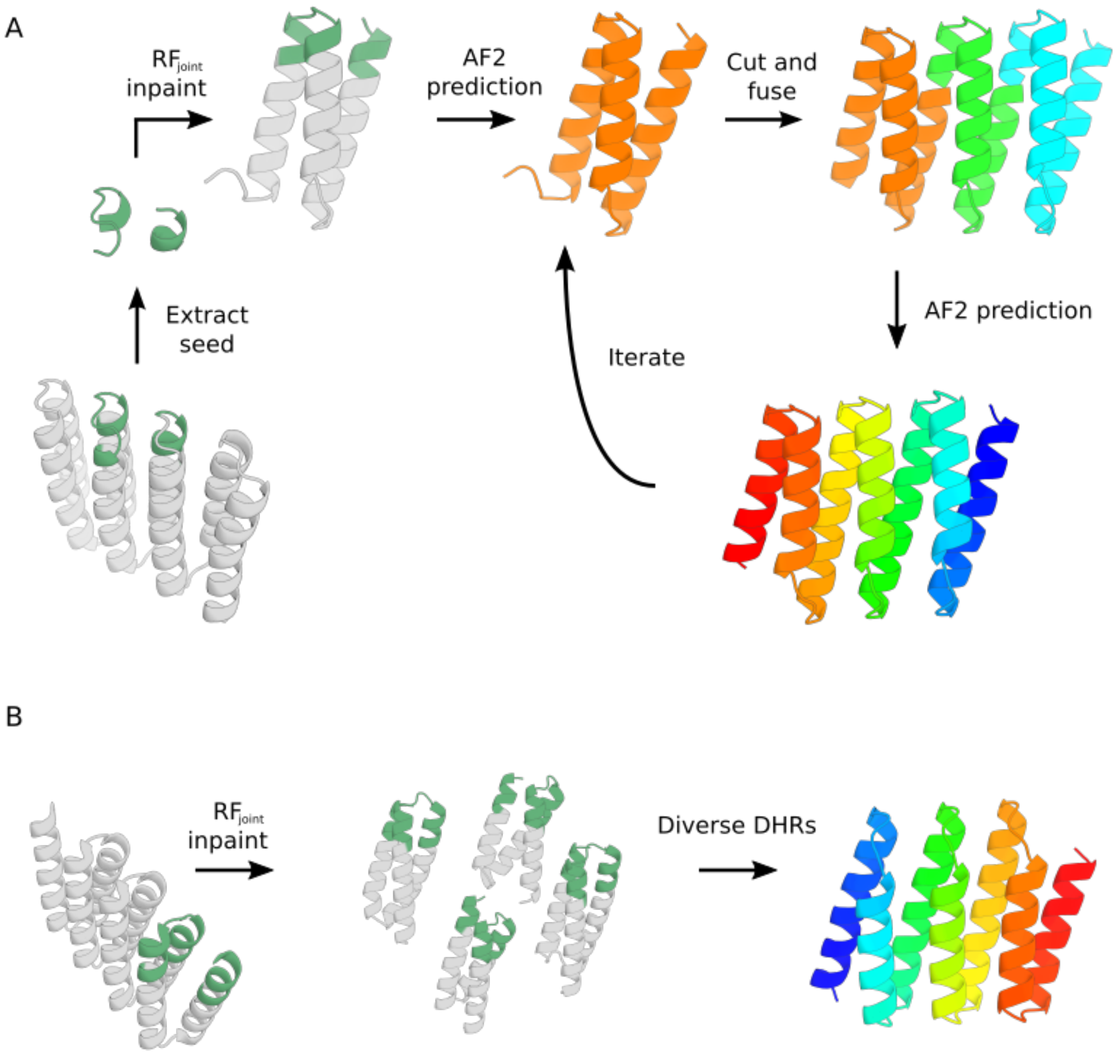


**Figure S20. Generation of designed helical repeat proteins (DHRs) with RF*joint***

Protocol for generation of designed helical repeat proteins (DHRs) by providing a native seed (green) as input. (A) Seed is chosen to provide information about designed symmetry, and an ensemble of designs are then generated. The designs are then aligned to the reference, symmetrically propagated, and fused. Design shown has AF plDDT 87 and backbone RMSD between RF and AF models of 1.6 Å when folded as a repeat with AF(B) Diversity is achieved by generating multiple RF*joint* designs and fusing samples with high lDDT scores. Example design has plDDT 87 and RMSD 1.3 Å when folded as a repeat with AF.

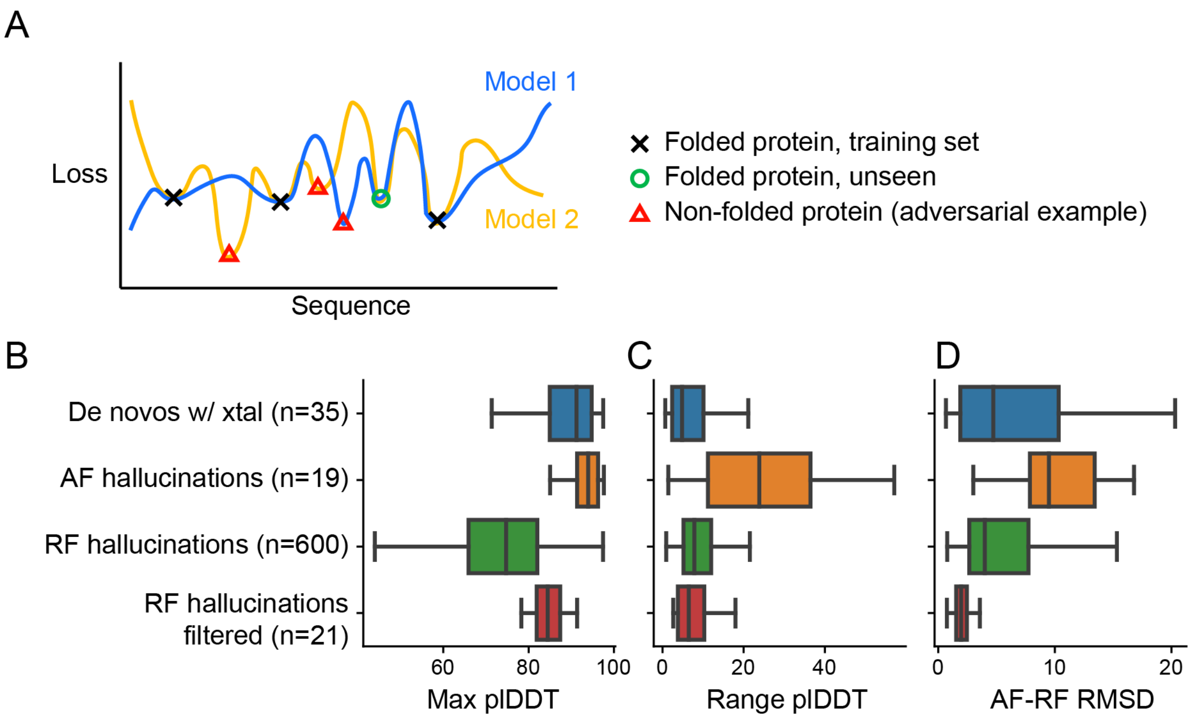


**Figure S21. Variability in neural net predictions for real and hallucinated proteins**

(A) Schematic of the loss landscape of 2 hypothetical neural networks independently trained on some data (X's, e.g. PDB protein structures). The models may have artifactual minima, where losses are low but the sequences do not correspond to folded and/or functional proteins (triangles). These "adversarial examples" may be sampled by methods like hallucination that seek to locally minimize the loss. Our hypothesis is that independently trained models should have similar losses for biophysically reasonable proteins, including those not seen during training (circles), but disagree with each other at inputs that do not fold well. To test this, we compared RoseTTAFold (RF) and AF predictions for: (1) de novo proteins with known crystal structures which were not in either RF or AF training sets, (2) hallucinations made by optimizing against AF model 5 (Supplementary Text), (3) hallucinations made with RF (random subsample of all hallucinations shown in Fig. S7), and (4) RF hallucinations after filtering on AF plDDT and motif RMSD and manual visual inspection (designs shown in Fig 2, 3, and S8). The AF hallucinations have the highest max predicted lDDT (B) across the 5 published AF models, but also the widest plDDT range (max - min) across the AF models (C). By contrast, plDDT range is lowest for the de novo proteins with known structures and almost as low for the RF hallucinations, suggesting that our pipeline may be more robust to adversarial example generation than methods that optimize against a single model. (D) Same comparison groups as (C) but plotting the RMSD between RF and AF (model 4) predictions. Note that the filtered RF hallucinations are expected to have low RMSD here due to the selection procedure (Methods).

### Supplementary Tables

**Table S1. Natural proteins used for mimetic design**

"Motif residues" indicate residues that were constrained to native geometry during hallucination. Semicolons delimit multiple sets of motif residues that were tested for the same problem. Sometimes only a subset of the motif residues actually comprise a binding interface or catalytic site; these are denoted "functional residues".

| Native protein (Reference) | PDB ID | Chain | Motif residues | Functional residues | Binding partner(s) |
|---|---|---|---|---|---|
| HAC PD-1 (*67*) | 5IUS | A | A63-82, A119-140 | A64, 66, 68, 70, 73-75, 77-78, 81, 85, 89-91, 124, 126, 128, 132, 134, 136, 139 | PD-L1 |
| RSV-F site II (*68*) | 3IXT | P | P254-277 | | Antibody |
| RSV-F site V (*19*) | 5TPN | A | A163-181 | | Antibody |
| ACE2 (*69*) | 6VW1 | A | A24-42 | | SARS-CoV2 receptor binding domain |
| EF-hand (*70*) | 1PRW | A | A21-31,A56-67 | A21-31,A56-67 | $Ca^{2+}$ |
| Di-Fe (*21*) | 1BCF | A | A18-25,A47-54,A94-97,A123-130 | A18, 51, 54, 94, 127, 130 | $Fe^{2+}$ |
| Carbonic anhydrase II (*25*) | 5YUI | A | A62-65,A93-97,A118-120 | A94,A96,A119,A199 | $Zn^{2+}$ |
| $\Delta^5$-3-ketosteroid isomerase (*28*) | 1QJG | A | A14,A38,A99 | A14,A38,A99 | equilenin |
| C3d (*71*) | 1GHQ | A | A104-126,A170-185; A109-117; A170-185 | | Complement receptor 2 (CR2) |
| B7-2 (*72*) | 1I85 | B | B84-88 | | CTLA-4 |
| IL-10 | 1Y6K | L | L34-40 | | IL-10R |

**Table S2. RMSDs between native protein, design model, and AlphaFold model**

All RMSDs are in angstroms. Columns in red are the metrics reported in the main text and figures.

| | Overall | | Motif | | |
|---|---|---|---|---|---|
| **Design** | **AF plDDT** | **RMSD, Design to AF** | **RMSD, Design to AF** | **RMSD, Design to native** | **RMSD, AF to native** |
| rsvf-ii_141 | 85 | 1.25 | 0.66 | 0.69 | 0.53 |
| rsvf-ii_158 | 83 | 1.37 | 0.74 | 0.93 | 0.51 |
| rsvf-ii_171 | 88 | 2.32 | 0.63 | 0.57 | 0.69 |
| rsvf-ii_118 | 78 | 0.96 | 0.63 | 0.51 | 0.69 |
| rsvf-ii_29 | 85 | 1.30 | 0.62 | 0.60 | 0.93 |
| rsvf-v_854 | 82 | 2.45 | 0.65 | 0.71 | 0.75 |
| rsvf-v_870 | 80 | 1.37 | 0.41 | 0.70 | 0.76 |
| rsvf-v_828 | 78 | 1.40 | 0.43 | 1.22 | 0.59 |
| rsvf-v_903 | 88 | 1.75 | 0.66 | 1.67 | 0.71 |
| rsvf-v_1050 | 75 | 1.82 | 0.63 | 0.73 | 0.74 |
| ace2_76 | 89 | 0.91 | 0.43 | 0.47 | 0.55 |
| ace2_1157 | 80 | 1.22 | 0.31 | 0.56 | 0.47 |
| ace2_1007 | 83 | 2.21 | 0.43 | 0.33 | 0.57 |
| ace2_1846 | 84 | 0.96 | 0.34 | 0.50 | 0.69 |
| ace2_600 | 80 | 1.98 | 0.48 | 0.56 | 0.70 |
| ace2_109 | 81 | 1.94 | 0.23 | 0.45 | 0.52 |
| efhnd_1m62 | 80 | 1.11 | 0.44 | 0.37 | 0.37 |
| efhnd_2m81 | 84 | 1.33 | 0.69 | 0.45 | 0.87 |
| dife_86 | 84 | 1.97 | 0.89 | 0.40 | 0.90 |
| dife_56 | 84 | 2.28 | 0.74 | 0.46 | 0.87 |
| dife_38 | 79 | 1.40 | 0.81 | 0.53 | 0.92 |

| | | | | | |
|---|---|---|---|---|---|
| dife_103 | 83 | 1.51 | 0.77 | 0.49 | 0.96 |
| dife_92 | 79 | 1.40 | 0.81 | 0.37 | 0.93 |
| hca_1 | 73 | 1.44 | 0.73 | 0.75 | 1.04 |
| hca_2 | 71 | 1.62 | 0.46 | 0.46 | 0.62 |
| ksi_1 (AF hallucination) | 84 | 1.04 | 0.30 (Cb) | 0.30 (Cb) | 0.30 (Cb) |
| ksi_2 (AF hallucination) | 72 | 1.06 | 0.16 (Cb) | 0.43 (Cb) | 0.53 (Cb) |
| IL10_179 | 82 | 1.40 | 0.21 | 0.22 | 0.35 |
| IL10_65 | 88 | 1.54 | 0.16 | 0.36 | 0.37 |
| IL10_71 | 75 | 1.56 | 0.40 | 0.29 | 0.45 |
| C3D_45 | 81 | 1.50 | 0.59 | 0.66 | 0.71 |
| C3D_79 | 70 | 2.02 | 0.28 | 0.24 | 0.28 |
| C3D_58 | 86 | 1.35 | 0.94 | 1.25 | 0.47 |
| B72_10 | 81 | 0.67 | 0.22 | 0.22 | 0.29 |
| B72_5 | 87 | 1.31 | 0.11 | 0.21 | 0.23 |
| B72_3 | 81 | 1.50 | 0.19 | 0.25 | 0.25 |

**Table S3. Frequency of suitable native scaffolds**

| Native protein | PDB ID | Chain | Motif residues | Scaffolds in the PDB with <1Å motif RMSD | |
|---|---|---|---|---|---|
| | | | | **Number** | **Frequency** |
| RSV-F site II | 3IXT | P | P254-277 | 0 | 0 |
| RSV-F site V | 5TPN | A | A163-181 | 1 | 3.76e-05 |
| ACE2 | 6VW1 | A | A24-42 | 1874 | 7.05e-02 |
| EF-hand (double) | 1PRW | A | A21-31,A56-67 | 30 | 1.13e-03 |
| EF-hand (single) | 1PRW | A | A56-67 | 77 | 2.90e-03 |
| Di-iron | 1BCF | A | A18-25,A47-54,A94-97,A123-130 | 3 | 1.13e-04 |
| Carbonic anhydrase II | 5YUI | A | A62-65,A93-97,A118-120 | 1 | 3.76e-05 |
| C3d | 1GHQ | A | A104-126,A170-185 | 2 | 7.52e-05 |
| HAC PD-1 | 5IUS | A | A63-82, A119-140 | 56 | 2.11e-03 |

### Supplementary Algorithms

##### Algorithm S1. Joint RosettaFold training Epoch

**Algorithm 1:** $RF_{joint}$ training epoch (pseudocode)

```
Data: D = (X_1, X_2, ..., X_n), X_i ∈ PDB
Batch size: B = 16

# enumerate through all PDB examples
for (i, X_i) in enumerate(D) do

    # randomly choose task
    n ← np.random.uniform(low=0.0, high=1.0)

    if n < 0.25 then
        # prepare structure prediction example
        X ← prepareInput(X, structure=True)

    else
        # prepare fixed BB sequence design example
        X ← prepareInput(X, structure=False)

    # input partial seq/str information
    # output completed seq/str
    Y ← RF_joint(X)

    # calculate loss, backward pass to accumulate gradient
    J ← calculateLosses(Y)
    J.backward()

    # step down gradient if batch is complete
    if (i modulus B) == 0 then
        optimizer.step()
    end
end
```

### Supplementary Data

##### Data S1. PDB files of all designs shown in paper
